# Capturing learning on the fly: an eye-tracking method to quantify prediction errors and updating the prior

**DOI:** 10.64898/2026.03.09.710486

**Authors:** Flóra Hann, Cintia Anna Nagy, Zita Olivia Nagy, Dezso Nemeth, Orsolya Pesthy

**Author notes:** corresponding authors: Flóra Hann, ELTE Eötvös Loránd University, 1071 Budapest, Damjanich utca 41-43., Hungary;, Orsolya Pesthy, CRNL - INSERM - U1028, 95 Bd. Pinel, Bâtiment 452, CH LE Vinatier, 69500 Bron, France. These authors contributed equally to this work (Flóra Hann, Cintia Anna Nagy). These authors contributed equally to this work (Dezso Nemeth, Orsolya Pesthy). senior author (Dezso Nemeth).

## Abstract

The ability to build predictive models of the environment fundamentally drives adaptive behavior. Yet, the real-time dynamics of how these internal models are formed and updated remain poorly understood. Conventional methods often rely on indirect, offline measures or noisy motor responses, limiting insight into the fine-grained computational processes underlying learning. Here, we introduce a generalizable, gaze-based analytical framework that directly tracks the trial-by-trial dynamics of expectation formation and updating. Applying this framework to an unsupervised probabilistic learning task, we categorized anticipatory saccades to dissociate prediction errors arising from environmental stochasticity from those reflecting an inaccurate internal model, and quantified how these predictions were iteratively revised. Learners differentiated between these error types: noise-driven errors were more likely to happen, and triggered less updates than errors reflecting insufficient knowledge of the regularity. At the same time, participants exhibited a strong preference to repeat their previous predictions. This repetition bias was amplified when predictions aligned with the underlying regularity, but was also present for non-aligned responses. Critically, updating depended more strongly on whether a prior belief was consistent with the task’s probabilistic structure than on whether the predicted stimulus matched the actual, presented stimulus. These findings suggest that statistical learning may not strongly be driven by errors; rather, it may rely on conservative updating with relatively low learning rate, or, on a Hebbian, repetition-based process. Our framework thus offers a dual contribution: a broadly applicable tool for quantifying real-time expectations, and evidence for a learning strategy that prioritizes model stability in noisy environments.

## Introduction

Predictive processing has been one of the most widely researched areas of cognitive science (Clark, 2013, 2016; Hohwy, 2013; Miłkowski & Litwin, 2022). It is thought to be fundamental to most cognitive processes, including perception and learning, which play a key role in adapting to our environment (Clark, 2013; Gregory, 1980). These processes require the brain to construct internal models of the environment using two sources of information: prior experiences (*priors*) and current sensory inputs – this is the mechanism referred to as predictive processing (Bastos et al., 2012; Clark, 2013; Friston, 2010; Keller & Mrsic-Flogel, 2018; Rao & Ballard, 1999). Notably, inputs from real-world environments are often ambiguous, and the regularities that we can use to build internal models are embedded in noise (Kelly et al., 2018; Williams, 2018). Statistical learning, a hallmark of predictive processing, is the mechanism the brain uses to extract these regularities from among elements of noise – often implicitly (Aslin, 2017; Howard & Howard, 1997). This ability is crucial to several everyday behaviors involving, e.g., social, linguistic, and musical skills (Lieberman, 2000; Romano Bergstrom et al., 2012; Ullman, 2016). As such, statistical learning has been a central topic of cognitive research for decades (Aslin, 2017; Frost et al., 2019), however, its measurement still poses some challenges. Many tasks use post-exposure (offline) tests that cannot grasp real-time learning (Lukács et al., 2021; Siegelman et al., 2018), rely on noisy manual responses (see e.g., Howard & Howard, 1997; Nissen & Bullemer, 1987; Schlichting et al., 2017) and thus fail to disentangle underlying mechanisms, meaning that performance does not exclusively reflect the process in question (see Németh et al., 2013). For example, most tasks do not allow for the precise separation of prior expectations (e.g., the implicit knowledge about what the next stimulus will be in a sequence learning task) from stimulus-response mappings (e.g., which button to press when a stimulus appears in a specific location). There is substantial evidence, however, that eye movements offer a more direct and sensitive measure of statistical learning than motor responses (Tal et al., 2021; Tal & Vakil, 2020; Vakil et al., 2017; Zolnai et al., 2022), as they capture anticipatory behavior – a proxy for prior expectations (Vakil et al., 2017). In this study, we aimed to overcome limitations and leverage eye movements to advance a more mechanistic understanding of statistical learning. To this end, we developed a novel methodological framework to track learning trajectories on the fly and to examine the role of prior expectations in statistical learning.

Eye-tracking has been increasingly used to study statistical learning, as anticipatory eye movements can provide insight into how predictions evolve over time. Early work demonstrated that infants’ gaze patterns during cross-situational word learning can be explained by associative statistical learning models – representing one of the first examples of the use of eye tracking to infer underlying processes in statistical learning (Yu et al., 2012; Yu & Smith, 2011). Subsequent studies extended this approach to adult learning tasks, demonstrating that anticipatory gaze shifts during serial reaction time tasks reveal sequence knowledge, often more sensitively than manual reaction times (Compostella et al., 2025; Tal et al., 2021; Vakil et al., 2017; Zolnai et al., 2022). In visual search settings, oculomotor measures like micro-saccades in pre-stimulus intervals reveal statistically learned suppression of distractors (Chen et al., 2025). Despite this rich literature, the notion that anticipatory eye movements directly reflect *priors* (which are central elements of predictive processing) has not been shown explicitly. Aiming to fill this gap, we further extend the new approach of Zolnai et al. (2022), who developed an oculomotor version of a reliable and widely used statistical learning task (the Alternating Serial Reaction Time, ASRT task; Farkas et al., 2024; Howard & Howard, 1997), which replaces manual responses with gaze tracking. Beyond offering a direct window to underlying mechanisms, requiring no button presses minimizes inevitable noise stemming from the motor system, ensuring a more process-pure measurement. However, former approaches to measure statistical learning by eye-tracking relied on anticipatory eye movements utilizing large Areas of Interest (AOI) around the target. While useful, this approach lacks the spatial resolution to distinguish fine-grained predictive dynamics. To overcome this, we employed saccades, that is, rapid, voluntary eye movements that shift the gaze from one point of interest to another (Dodge, 1903; Westheimer, 1954). Building on the probabilistic structure of the ASRT task and more fine-grained eye-tracking measurements, here, we introduce novel indices of priors and their updating. In doing so, we aim to lay the groundwork for metrics that can be used in the context of any probabilistic task and will thus advance the study of statistical learning, particularly the role of predictions.

Accumulating evidence indicates that statistical learning is not a unified concept (Simor et al., 2019; Szegedi-Hallgató et al., 2017; Takacs et al., 2024; Vékony et al., 2023), and anticipatory eye movements may reflect these multiple underlying mechanisms (Németh et al., 2026; Németh & Tóth-Fáber, 2026; Tal et al., 2021). How these different mechanisms contribute to probabilistic statistical learning, however, remains poorly understood. Investigating this question requires a task capable of distinguishing errors that reflect a lack of knowledge from those that arise from noise. A probabilistic task structure allows precisely this distinction between anticipatory errors. Here, we demonstrate this framework on the eye tracking version of the ASRT task (Zolnai et al., 2022), a relatively reliable (Farkas et al., 2024) implicit task (Vékony, Ambrus, et al., 2022) that is more process-pure compared to many other statistical learning paradigms. However, our approach is general and can be adapted to any statistical learning task involving probabilistic stimulus streams. Because the stimuli follow an alternating probabilistic sequence, upcoming events can never be predicted with full certainty: some stimuli appear with a higher, some with a lower probability. As a result, anticipatory errors may occur even when the response corresponds to the underlying regularity. We term these *learning-dependent errors* and distinguish them from *not-learning-dependent errors*, which reflect an inaccurate representation of regularities (see Table 2 in detail). With eye-tracking, we can register anticipatory saccades, that is, the first eye movement after the previous stimulus disappeared. Note that since anticipatory eye movements are recorded in the absence of a stimulus, they reflect relatively pure expectations independent of external stimuli. In this study, we present a detailed methodology for measuring learning-dependent and not-learning-dependent errors via eye-tracking, and we analyze the dynamics of these metrics across the task. As such, we aim to provide a tool for the fine-grained investigation of expectation formation during learning – adaptable to any probabilistic learning paradigm.

In addition to examining errors related to the predictive process, understanding how prior expectations are updated over time is crucial for shedding light on the dynamics of statistical learning. In our study, we capture the iterative updating of priors by assessing whether participants change their anticipatory gaze from one occurrence of a predictive unit to the next. This approach allows us to track whether a prior is retained or modified, reflecting the ongoing adjustment of internal models. Moreover, to better understand precisely what these updates reflect, we further categorize updates based on whether they represent learning-dependent changes – shifts toward high-probability stimuli – or not-learning dependent changes, which represent gaze shifts toward low-probability (uninformative) locations. This distinction enables us to infer when participants are consolidating or disrupting accurate predictions, correcting task-incongruent internal models, or engaging in exploratory behavior. Importantly, the method provides a fine-grained, trial-by-trial measure of learning dynamics, offering novel insights into how priors are formed, maintained, and revised over time – a central aspect of understanding predictive processing (Clark, 2013; Friston, 2005, 2010).

Our study had two main aims. First, we sought to develop a gaze-based measure of the learning process that can be applied to any learning task involving probabilistic stimulus streams. Building on the framework of learning-dependent and not-learning-dependent eye movements established by Zolnai et al. (2022), we further distinguished between correct and erroneous trials, utilizing the fact that in probabilistic tasks, correct responses are not always learning-dependent. Second, we aimed to capture how priors are updated over the course of learning. To this end, we developed metrics to track how prior expectations are updated iteratively on a trial-by-trial basis, differentiating updates that consolidate accurate predictions from those reflecting noise, exploration, or a lack of knowledge of the regularity. This framework leads to specific, falsifiable predictions in line with our aims. First, as participants learn the statistical structure, we expect anticipatory errors to increasingly align with the underlying regularities; consequently, we expect the likelihood of learning-dependent errors to progressively increase as compared to not-learning-dependent errors. Second, we expect that if our iterative update measure indeed captures the adjustment of prior expectations, then (a) not-learning-dependent saccades will be updated more frequently than learning-dependent ones; and (b) these updates will increasingly converge on high-probability locations, reflecting a refinement of the internal model.

## Methods

### Participants

A total of 185 healthy young adults with normal or corrected-to-normal vision were recruited through a university course from Eötvös Loránd University in Budapest and received course credits for participation. In line with the inclusion criteria, none of the participants reported a history of neurological impairment or head injury, a diagnosis of any psychiatric condition or epilepsy, or having consumed alcohol within 6 hours, recreational drugs or psychoactive medications within 24 hours prior to the experiment.

Due to the unsuccessful calibration of the eye-tracker, 27 participants were excluded. We excluded an additional 5 participants due to software issues or experimenter error. Thus, we used the data of 153 participants (115 females (75.16%), 38 males (24.84%); M_age_ = 22.2 years ± 5.69 SD, range = 18–53 years). Additionally, 25 participants were excluded from the analyses following outlier filtering for eye-tracking data quality (see *Quality control of data* section). Our final sample therefore consisted of 128 participants (96 females (75%), 32 males (25%); M_age_ = 22 years ± 5.64 SD, range = 18–53 years).

All participants gave informed consent, and the study was approved by the Regional and Institutional Committee of Science and Research Ethics, Semmelweis University, Budapest, Hungary (SERKEB No.: BM/6621-1/2024). The experiment took place at the laboratory of the Brain, Memory and Language Lab, Eötvös Loránd University, Budapest.

### Task & Procedure

To assess statistical learning, we employed an eye-tracking version of the Alternating Serial Reaction Time (ASRT) task (Zolnai et al., 2022). Data was recorded using the Tobii Pro Python SDK (Tobii AB, 2020b) integrated with a PsychoPy-based (Peirce et al., 2019) experimental script. The task script used in this study is available on GitHub (Project ET Developers, 2020/2021). In the task, participants were exposed to a probabilistic sequence. They viewed four empty circles arranged in a square in each corner of the screen, with one circle turning blue on each trial, indicating the activation of the stimulus (Figure 1A). Participants were instructed to fixate on the blue circle as quickly as possible. The specific instructions were as follows: ‘*You will see four empty circles on the screen, one of which will turn blue from time to time. Your task is to move your gaze to the location of the circle that has turned blue. Follow the appearing stimuli with your eyes as accurately and quickly as possible! The blue circle may appear in the same place several times in a row.*’. Once they fixated on the target for at least 100 ms, the blue stimulus activation disappeared, and the next circle turned blue after a 500-ms response-stimulus interval (RSI). We refer to oculomotor reaction time (oRT) as the time from stimulus onset to the beginning of the 100 ms fixation on the target stimulus (Figure 1B). The circles had a 3 cm diameter (≈ 2.64° visual angle) and they were spaced at an equal distance from each other (15 cm, ≈ 13.16° visual angle) and from the center of the screen (10.6 cm, ≈ 9.32° visual angle).

**Figure 1.**
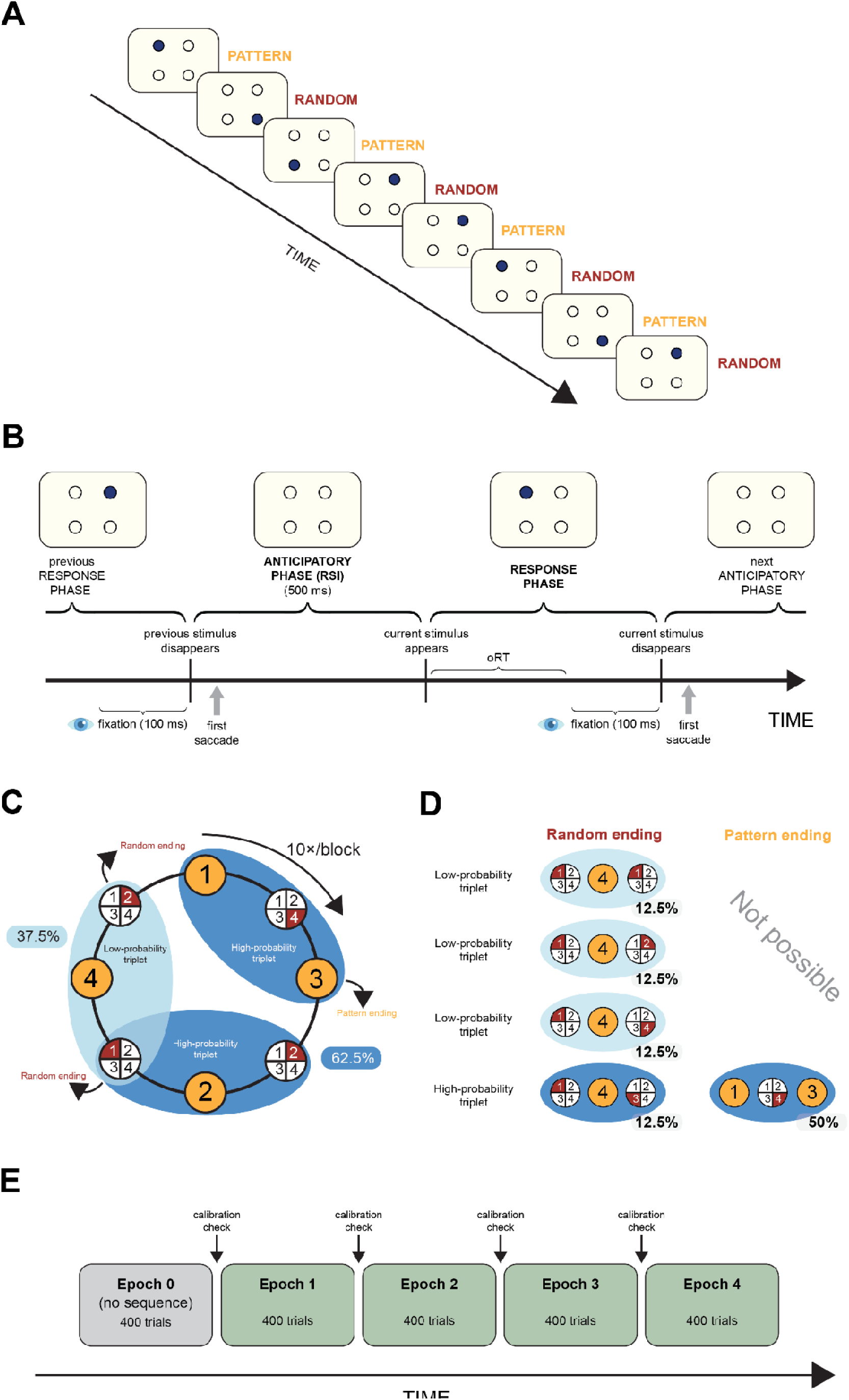
Experimental design and structure of the oculomotor ASRT task. **A)** In the task, participants viewed four empty circles arranged in a square in each corner of the screen, with one circle turning blue on each trial. In the stream of stimuli, every second trial was part of an 8-element probabilistic sequence. Random elements were inserted among pattern elements to form the sequence (e.g., 1-R-3-R-2-R-4-R, where numbers indicate the location of one of the four circles on the screen, and R represents random positions). **B)** Each trial consisted of two phases: an anticipatory phase (response–stimulus interval, RSI) and a response phase. On each trial, one of four empty circles turned blue, signaling the target stimulus. Participants were instructed to shift their gaze as quickly and accurately as possible to the blue circle and maintain fixation for at least 100 ms. Once fixation was detected, the stimulus disappeared, and after a 500-ms RSI, the next target appeared. Oculomotor reaction time (oRT) was defined as the latency from target onset to fixation on the target location. The schematic illustrates the temporal sequence of events: disappearance of the previous stimulus, the anticipatory phase, onset of the current stimulus, response phase, and transition to the next anticipatory phase. **C)** Formation of triplets in the task. Pattern elements are represented by orange backgrounds (they appear consistently in the same position throughout the task), and random elements are represented by red backgrounds (they appear randomly in one of the four possible positions). Every trial can be identified as the third element of three consecutive trials (a triplet) in a sliding window manner. The probabilistic sequence structure results in some triplets occurring with a higher frequency (high-probability triplets, 62.5% of all trials, indicated with dark blue backgrounds on the figure) than others (low-probability triplets, 37.5% of all trials, indicated with light blue backgrounds on the figure). Statistical learning is operationalized as the performance improvement in high-probability trials compared to low-probability trials. Ten repetitions of the 8-element sequence (80 trials) make up one block of the task. **D)** The formation of high-probability triplets can involve the occurrence of either two pattern trials with one random trial between them, which occurs in 50% of trials; or two random trials with one pattern trial between them, which occurs in 12.5% of trials. In total, 62.5% of all trials constitute the final element of a high-probability triplet, while the remaining 37.5% are the final elements of a low-probability triplet. **E)** The task was organized into blocks, each consisting of 80 trials (ten repetitions of an 8-element sequence), preceded by five initial random warm-up trials that were excluded from all analyses. A unit of 5 blocks is referred to as an epoch, which thereby contains 400 trials. Participants were familiarized with the task by completing one practice epoch containing only random trials. The actual task comprised four epochs containing the alternating sequence. After every epoch, calibration was reassessed based on the last 20 trials, and recalibration was performed if necessary.

Unbeknownst to the participants, the sequence of stimuli followed a hidden pattern. Specifically, the appearance of the stimuli followed a probabilistic eight-element sequence where pattern and random elements alternated with each other (Figure 1A; e.g., 1-R-3-R-2-R-4-R, where the numbers represent one of the four circles on the screen, and “R” indicates a randomly selected circle). This probabilistic structure resulted in certain combinations of three consecutive stimuli (triplets) being more probable to appear than others: high-probability triplets followed the underlying sequence, arising either from a pattern-random-pattern (P-R-P) or random-pattern-random (R-P-R) structure (e.g., 1-X-3, 3-X-2, 2-X-4, and 4-X-1 triplets, where “X” indicates either a pattern or random trial). In contrast, low-probability triplets were formed solely through the R-P-R combination and did not follow the underlying sequence (e.g., 1-X-2 or 1-X-4, where “X” indicates a pattern trial; Figure 1C). In total, high-probability triplets could be formed 16 different ways, while low-probability triplets could be formed 48 different ways. The individual probability of appearance was ∼3.91% for each high-probability triplet and ∼0.78% for each low-probability triplet, yielding a total probability of 62.5% (16 × 3.91) for high-probability, and 37.5% (48 × 0.78) for low-probability triplets (Figure 1D). When referring to “triplet type” throughout the rest of this paper, the focus is on trials serving as the last element of a high- or low-probability triplet.

There is robust evidence demonstrating performance facilitation in motor and oculomotor responses over time to high-probability triplets compared to low-probability triplets (Howard & Howard, 1997; Kóbor et al., 2017; Németh et al., 2010; Song et al., 2007; Takács et al., 2018; Tóth-Fáber et al., 2023). This indicates that participants become increasingly sensitive to the underlying probabilistic structure of the task. Furthermore, this knowledge remains entirely implicit, with self-reports and sequence-generation tasks finding no evidence of explicit awareness (Vékony, Ambrus, et al., 2022). To confirm this, we also asked our participants after the ASRT task whether they noticed anything unusual, specifically focusing on any regularities detected. None of the participants could give an accurate description of the underlying sequence or even noticed its existence. Therefore, the differences in performance between high- and low-probability triplets indicate implicit statistical learning.

The task was organized into blocks, each consisting of 80 trials (ten repetitions of an 8-element sequence), preceded by five initial random warm-up trials that were excluded from all analyses. After completing each block, participants were shown their average oRT for that block, allowing them to monitor and potentially improve their performance. They also had the option to take a short, self-paced break between blocks (<1 min). Participants were familiarized with the task by completing five practice blocks containing only random trials. These practice blocks were included in the analyses (except testing the update of prior knowledge; see in detail later) as they indicate the baseline to which performance can be compared. The actual task comprised 20 blocks containing the alternating sequence. As per the standard ASRT analysis, we refer to a unit of 5 blocks as an epoch (Figure 1E). Eye movements and fixations were recorded using the Tobii Pro Fusion eye-tracker with a 120 Hz sampling rate (Tobii AB, 2025). The device was mounted on the bottom of a 24-inch AOC LED monitor with a screen resolution of 1920×1080. An approximate subject-screen distance of 65 cm was maintained throughout the task. Further details on gaze position estimation and fixation identification are provided in Zolnai et al. (2022).

At the start of the task, eye-tracker calibration was performed using the Tobii Pro Eye Tracker Manager (Tobii AB, 2024). Dots appeared sequentially in each corner and the center of the screen. Participants were asked to look at each dot until it “exploded” and disappeared. Following this, calibration was validated using 20 random trials. If no extreme oculomotor reaction times (defined as oRTs over 1000 ms) were detected, the calibration was considered successful. If extreme oRTs were present, the calibration-validation process was repeated until no extreme responses were recorded. If calibration failed after six attempts, the experiment was terminated, and the participant was excluded. Additionally, after every fifth block (i.e., after every epoch), calibration was reassessed based on the last 20 trials, and recalibration was performed if necessary (Figure 1E).

### Quality control of data

To maintain the quality of eye-tracking data, we excluded participants based on data quality measurements similar to the ones used in Zolnai et al. (2022): precision, missing data ratio, and eye distance from the screen. Precision refers to the stability of the recorded gaze direction while the participant is fixating on the current stimulus. To assess this, we computed two types of precision scores. The root mean square of sample-to-sample differences (RMS(S2S)) measures the difference between consecutive gaze samples during a fixation – when it is assumed that the participant is looking at the same point. We calculated the median RMS(S2S) value for each epoch of each participant. However, due to a known upper limit in this measurement (discussed in detail in Zolnai et al., 2022), we included an additional precision metric: the root mean square of the eye-to-eye distance (RMS(E2E)). This metric captures the positional differences between the left and right eyes for each valid sample during a fixation (excluding any samples without valid data for both eyes) and calculates the RMS of these differences. Outlier epochs were identified using the boxplot method – specifically, values falling above the upper bound or below the lower bound of 1.5 times the interquartile range, as calculated based on RMS values in all epochs of all participants pooled together (153 participants × 5 epochs = 765 epochs in total). Participants were excluded if they were outliers in at least one epoch during the data collection. Additionally, we assessed data loss, which can result from factors such as blinking or eyelashes. For each participant, we determined the percentage of invalid samples per epoch. Participants with a missing data ratio of 20% or higher were excluded (based on existing protocols, e.g., Hessels et al., 2020). Lastly, we recorded each participant’s distance from the screen during the task. If the median distance in any of the epochs was outside of the device’s functional range (50-80 cm), the participant was excluded. The expected versus observed values for each measurement, along with the number of excluded epochs, are presented in Table 1. Code for data quality filtering is available on the following OSF repository: https://osf.io/a4xhn/.

**Table 1.**
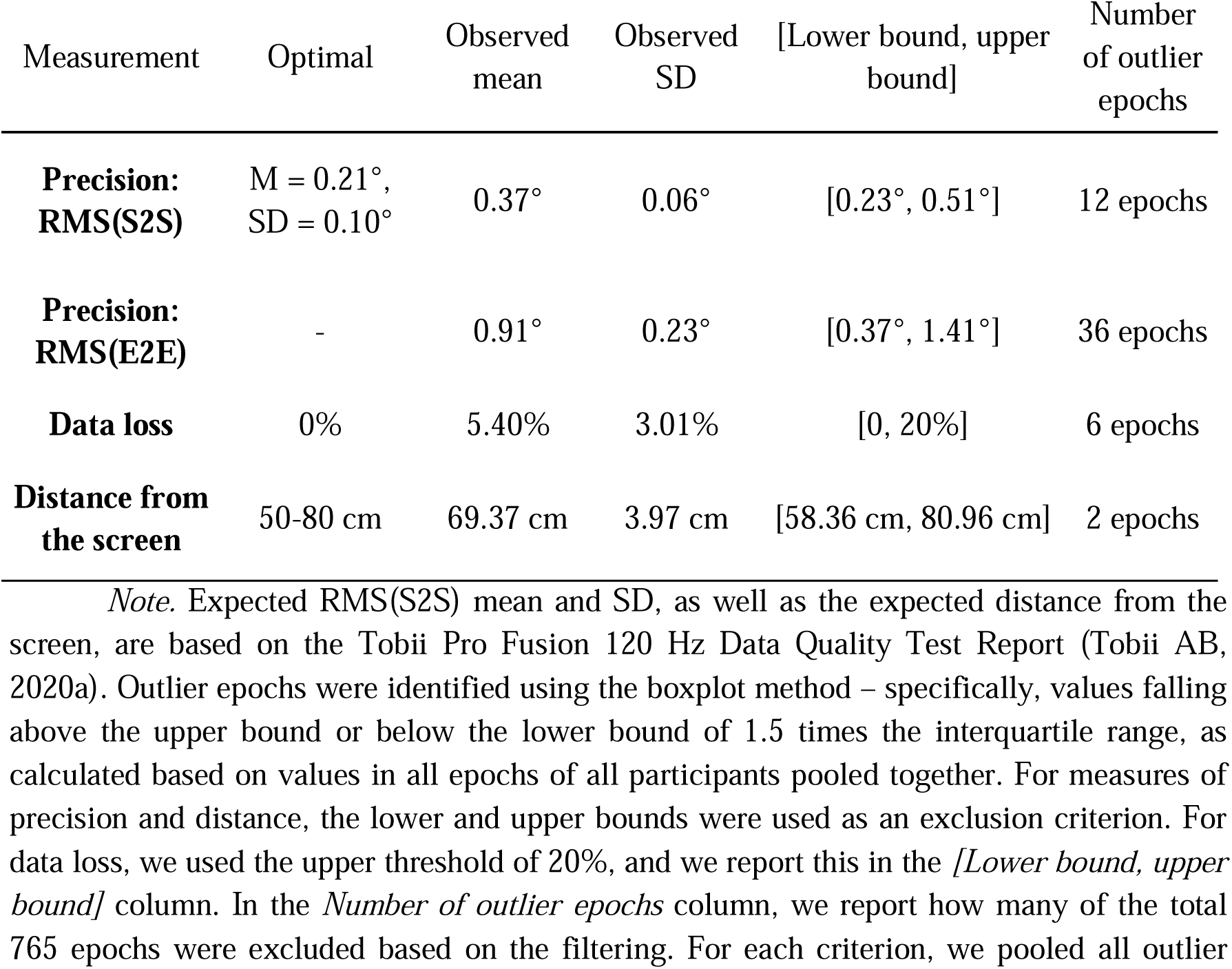
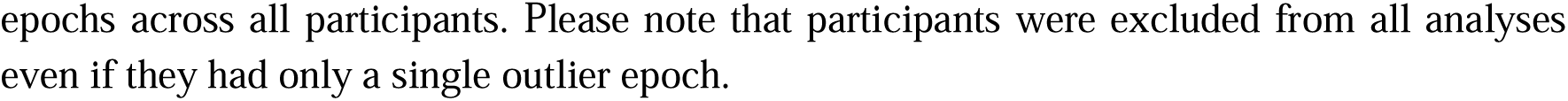
Data quality measures of eye tracking data.

### Data preprocessing and saccade extraction

Data were preprocessed in Python (version 3.13; Python Software Foundation, 2024), using the *NumPy* (version 2.4.1; Harris et al., 2020), *pandas* (version 3.0.0; McKinney, 2010; team, 2026), *SciPy* (version 1.13.1; Virtanen et al., 2020) packages. All preprocessing codes are available at https://osf.io/a4xhn/. We used a hybrid eye selection approach: left and right eye position coordinates were averaged where both were available, and, to minimize missing data, single eye position coordinates were used when the other eye position coordinates were unavailable. Samples where none were available were marked as missing.

For preprocessing oRT data, we used the same fixation identification algorithm as Zolnai et al. (2022). oRTs were defined as the time interval between stimulus onset and the onset of the first fixation on the corresponding stimulus location. A fixation was considered valid if the gaze remained within a 4x4 cm square around the stimulus location for at least 100 ms. For full algorithmic details, see Zolnai et al. (2022).

Saccade extraction followed a similar procedure to Lum (2020). To identify the onset and offset of all potential saccades, we computed the sample-to-sample eye velocity in °/s. We reduced high-frequency noise using a Savitsky-Golay filter (frame length: 7 samples, polynomial order: 5; Lum, 2020; Nyström & Holmqvist, 2010), via *savgol_filter* in the *scipy.signal* package. To ensure biological feasibility, a sample was classified as part of a saccade if its velocity exceeded 30 °/s but remained below 1000 °/s. In addition, we required that velocity within the onset-offset interval reached a minimum peak of 50 °/s, ensuring a clearly defined saccade profile. For each candidate saccade, duration (in ms) and amplitude (in degrees of visual angle) were computed. Saccades with durations shorter than 16 msec and longer than 100 ms, or amplitudes exceeding 30° were excluded to remove artifacts. For the descriptive statistics of saccades, see Supplementary Table S1.

For each retained saccade, the estimated saccade target was calculated and classified into one of the possible stimulus locations. Saccade vectors were computed in visual-angle space from gaze position at saccade onset to gaze position at the corresponding offset. For each saccade, vectors from the gaze position at onset to each stimulus location center were computed, and the angular difference between the saccade vector and each stimulus vector was calculated. The stimulus whose vector formed the smallest angle with the saccade vector (minimal angular deviation) was labeled as the predicted stimulus (see Figure 2). Saccades with minimal angular deviations of 90° or greater were excluded from further analyses, as these saccades indicated eye movements toward the screen edges, outside the task-relevant stimulus space (i.e., the four circles).

**Figure 2.**
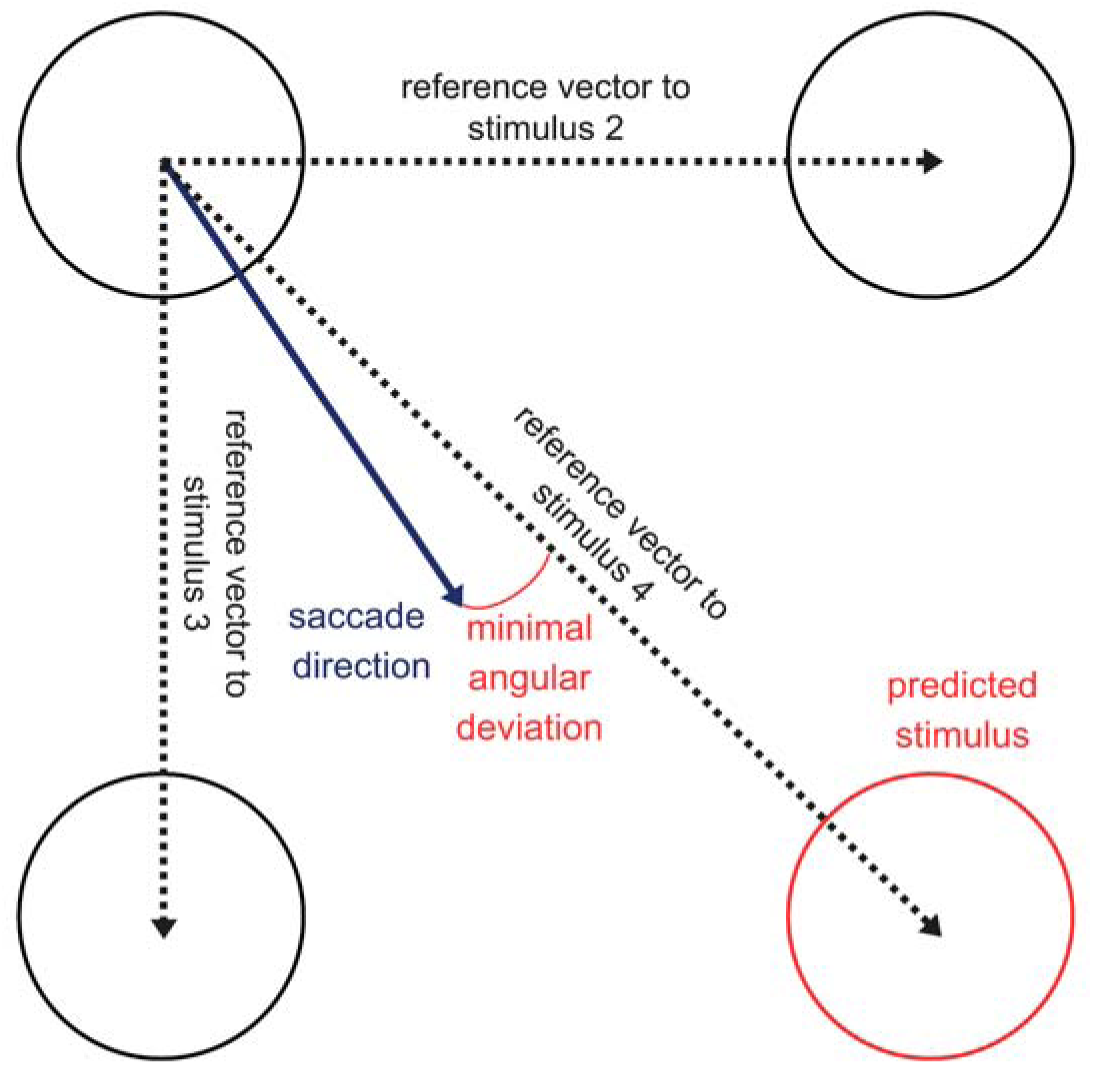
Determination of the predicted stimulus using minimal angular deviation. The schematic illustrates the method for classifying anticipatory saccades. The saccade vector (solid blue arrow) is defined by the gaze coordinates at saccade onset and offset. Reference vectors (dashed black arrows) connect the onset to the center of each of the potential stimulus locations. We computed the angular difference between the saccade vector and each reference vector; the stimulus location minimizing this deviation (minimal angular deviation; red arc) was identified as the predicted target stimulus.

Finally, we only retained saccades within each trial’s RSI, as in this period, due to the lack of sensory input, participants can only rely on their prior knowledge to guide their gaze. Where multiple saccades were present during the RSI, we considered only the first saccade, since this saccade is assumed to reflect early predictive influences shaped by prior experience during implicit learning (Jiang et al., 2014). Trials in which no such saccade was detected were marked as missing.

### Statistical analyses

#### General statistical modeling approach

Data analysis was performed in R (version 4.5.1; R Core Team, 2024). Linear and generalized linear mixed models were fitted with the *mixed* function of the *afex* package (version 1.4–1; Singmann et al., 2024). Models included Epoch coded as a factor to reflect the progression of the task in time, with the first sequential epoch as the reference in all models. Further, the specific variables of each model are detailed below in the subsections describing the models. Model syntaxes are also provided in general *lmer* notation. Each model included predictor variables as fixed effects and participant-specific intercepts as random effects. Where applicable, models included Epoch as a random slope. We tested assumptions of linearity, normal distribution of residuals, homoscedasticity, and multicollinearity using the *sjPlot* (version 2.9.0; Lüdecke, 2023), *performance* (version 0.15.0; Lüdecke et al., 2021), and *car* (version 3.1–1; Fox & Weisberg, 2019) package in R, evaluating scatterplots, QQ plots, and statistical tests. Assumptions of linearity and homoscedasticity were met in all cases. Recent work indicated that fixed effect estimates of linear mixed models are robust to violations of the assumption of residual normality (Knief & Forstmeier, 2021; Schielzeth et al., 2020). Thus deemed our models adequate, even though this assumption was slightly violated in all cases.

Summary tables (see Supplementary Materials) consistently report sample sizes both in terms of total data points and sampling units, along with random effects estimates, Nakagawa’s marginal and conditional R^2^s (Nakagawa & Schielzeth, 2013) and adjusted Intraclass Correlation Coefficients (ICCs). Post-hoc contrasts were carried out using the *emmeans* R package (version 1.11.2; Lenth, 2024) after regridding. Type III tests were used for inference on fixed effects, with degrees of freedom estimated via the Satterthwaite approximation for linear mixed models (Satterthwaite, 1941). For generalized linear mixed models, *p-values* were obtained via likelihood ratio tests. An alpha level of .05 was applied across all analyses. For multiple comparisons, *p* values from post-hoc tests were adjusted using the Šidák method (Šidák, 1967), and all *p* values are reported as two-tailed. Models were estimated with the *bobyqa* optimizer for linear mixed models and the *Nelder-Mead* optimizer for generalized linear mixed models, with optimizer settings adjusted to increase the maximum number of iterations (maxfun = 1e6) for the latter to facilitate convergence.

Figures were created using the *ggplot2* package (version 3.5.2; Wickham, 2016). Figures always plot raw data, whereas post-hoc tests are done using model-estimated marginal means. All data and codes for data analysis and visualization are available on the following OSF repository: https://osf.io/a4xhn/.

#### Metrics

We created a set of behavioral metrics to capture different aspects of statistical learning, anticipatory saccades, and updating processes. The name, abbreviation, operationalization, and interpretation of these metrics are presented in Table 2.

**Table 2.**
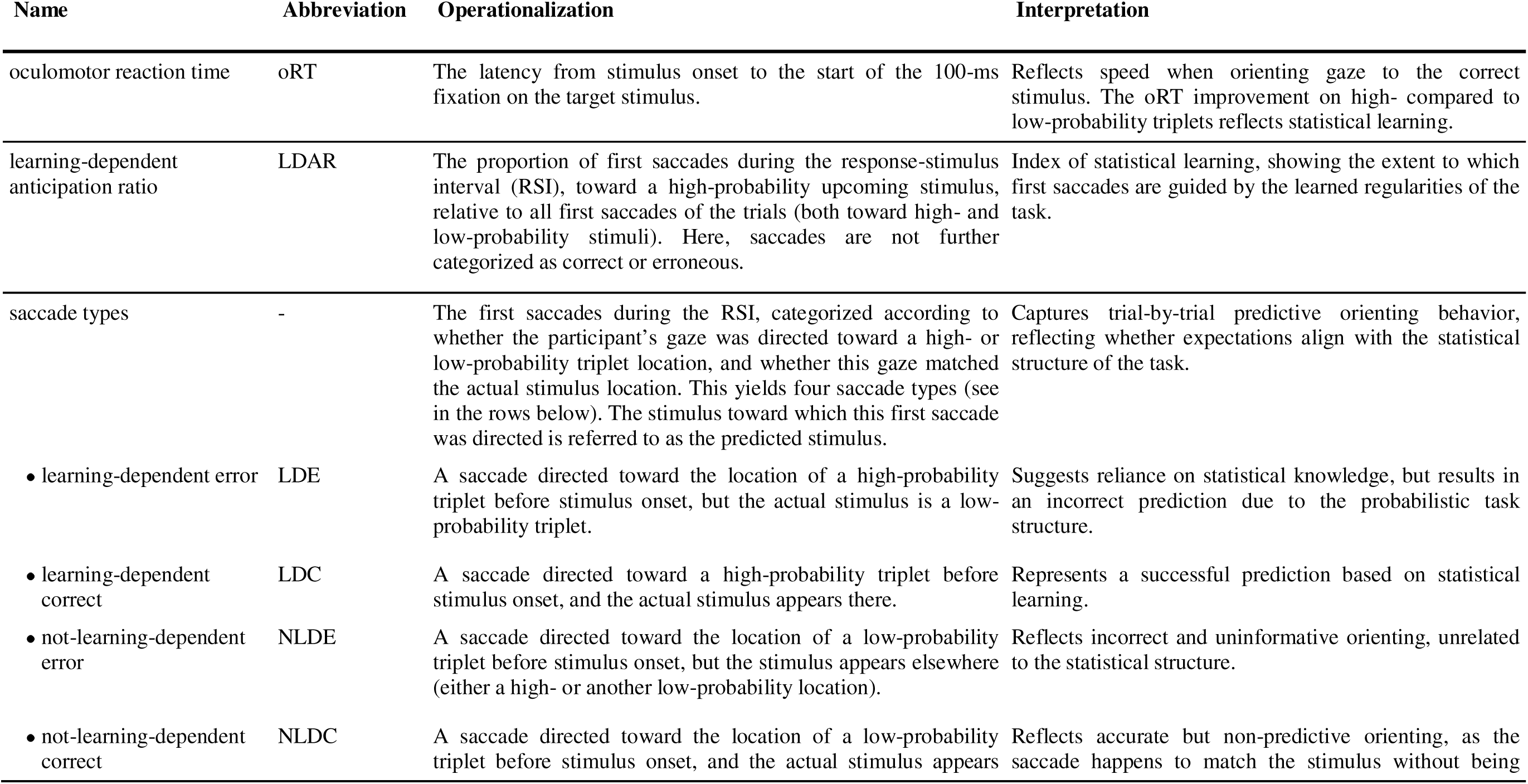
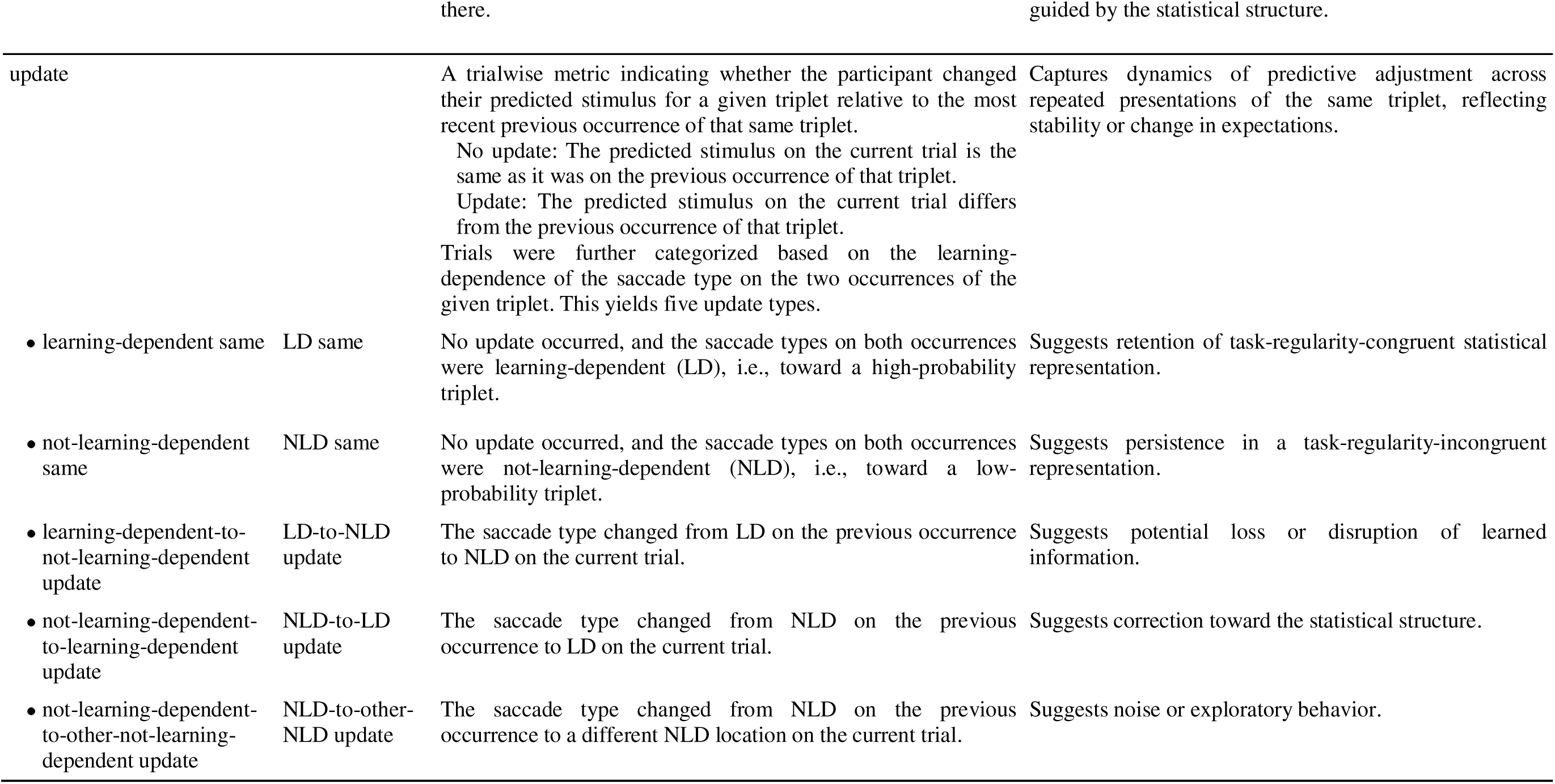
Metrics used for the analysis of eye tracking data.

#### Statistical learning – oculomotor reaction times and learning-dependent anticipation ratio

To test whether participants showed statistical learning, we applied the standard eye-tracking ASRT analysis protocol as described in Zolnai et al. (2022). First, we categorized each trial as high- or low-probability in a sliding window manner, based on whether it was the last element of a high- or a low-probability triplet. Trials that are the last elements of a repetition (e.g., 1-1-1) or a trill (e.g., 1-2-1) were removed, as participants tend to have a pre-existing tendency to react faster to these, unrelated to learning (Soetens et al., 2004). Please note that this exclusion happened only in the analysis of oRTs and learning-dependent anticipation ratios (see later).

When analyzing the oRT of fixations, we excluded trials with oRTs exceeding 1000 ms. Median oRTs were then computed separately for high- and low-probability triplets within each epoch for each participant. We fit a linear mixed model (Model_SL-oRT_) using our general modelling approach with epochwise median oculomotor reaction times as outcome variables, and Epoch (0-4) and Triplet type (high- or low-probability) as within-subject predictor variables.

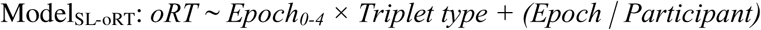

To assess whether participants made learning-dependent anticipations, we followed the logic of Zolnai et al. (2022), but instead of the last detectable gaze before stimulus appearance, we analysed the first saccades of the RSI. We compared the ratio of trials where this first saccade was directed toward a high-probability triplet (i.e., learning-dependent anticipation) compared to all trials where a saccade was made. This measure is henceforth referred to as the learning-dependent anticipation ratio (LDAR). We ran a linear mixed model (Model_LDAR_) using epochwise mean LDAR for each participant as an outcome variable, and Epoch (0-4) as a within-subject predictor variable.

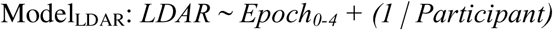

#### Likelihood of saccade types

In addition to the standard measures of the ASRT task, we applied a novel method. Each trial where a saccade was detectable, was categorized according to the participant’s first saccade direction during the RSI type. First, we distinguished learning-dependent (LD) and not-learning-dependent (NLD) saccade types, reflecting whether the participant moved their gaze toward a high- or a low-probability triplet, respectively (similarly to the analysis of LDAR). Each trial was then further classified as correct when the saccade was toward the stimulus location where the new stimulus actually appeared, or as an error when the saccade direction was toward a different location. This resulted in four final saccade types: ***learning-dependent correct*** (LDC, anticipated a high-probability triplet that actually appeared as the next stimulus), ***learning-dependent error*** (LDE, anticipated a high-probability triplet but a low-probability triplet appeared), ***not-learning-dependent correct*** (NLDC, anticipated a low-probability triplet that actually appeared), and ***not-learning-dependent error*** (NLDE, anticipated a low-probability triplet but another triplet, high- or low-probability, appeared; Figure 3).

**Figure 3.**
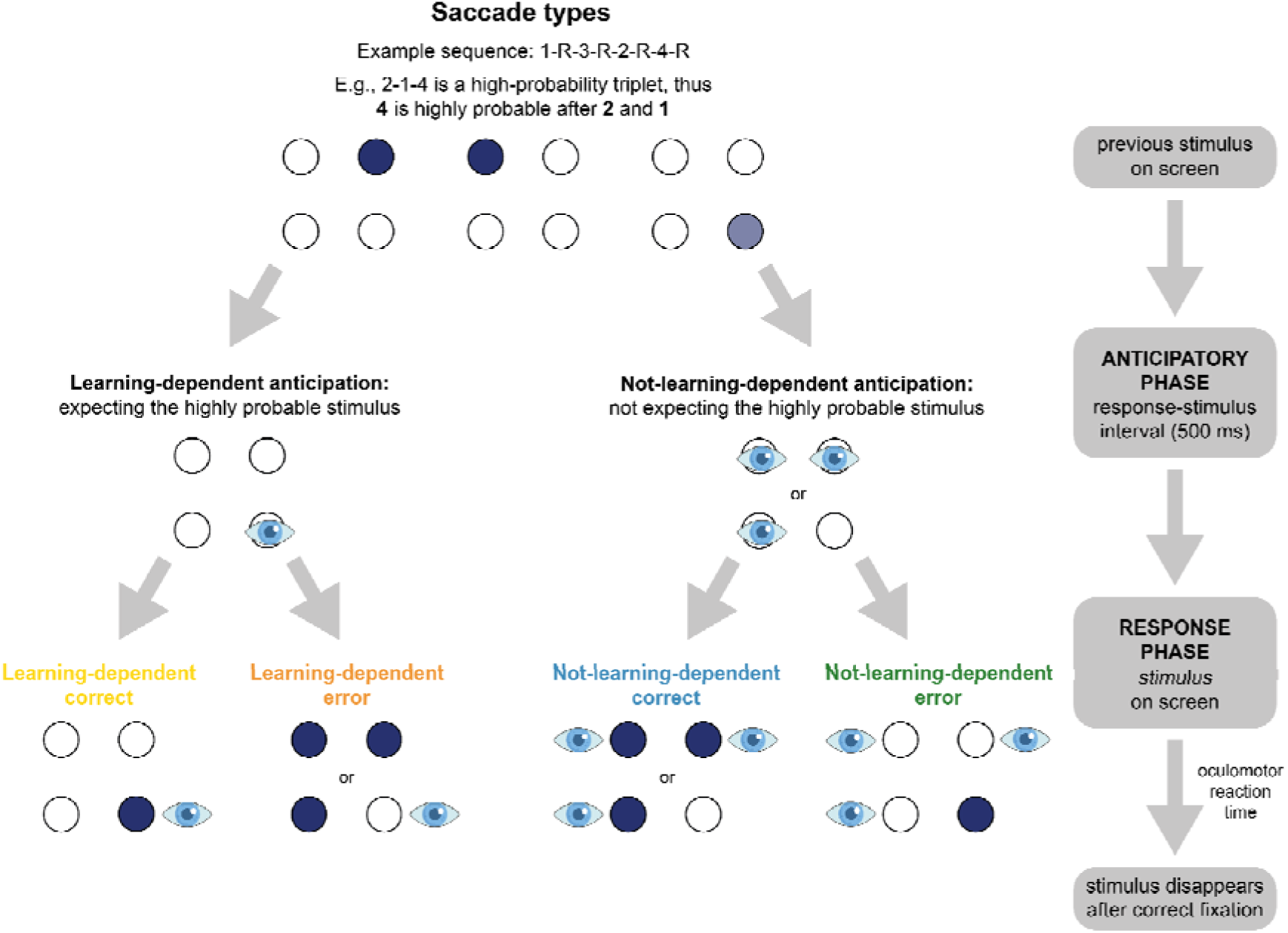
Categorizing saccades. During the anticipatory phase (response–stimulus interval, RSI), participants’ first valid saccade was recorded to capture anticipatory eye movements. Saccades toward a high-probability upcoming stimulus was defined as a learning-dependent (LD) anticipation, whereas saccades directed elsewhere were classified as not-learning-dependent (NLD). Each trial was further categorized as correct if the first saccade of the anticipatory phase was directed to where the stimulus subsequently appeared, or as an error if it was directed elsewhere. This resulted in four saccade categories: learning-dependent correct (LDC), learning-dependent error (LDE), not-learning-dependent correct (NLDC), and not-learning-dependent error (NLDE). The schematic illustrates the sequence of events, the detection of anticipatory gaze shifts, and their categorization into these four saccade types.

For each participant, we calculated the epochwise proportion of each saccade type. To ensure interpretability, the proportions were standardized against their chance-level probabilities. The baseline probability of these types is as follows: LDC: 0.15625, LDE: 0.09375, NLDC: 0.09375, NLDE: 0.65625; see Supplementary Table S2 for details. As a result, this metric reflects the likelihood of each trial type, with a chance-level of one. To analyze the likelihoods of saccade types and how they changed throughout the task, we fit a linear mixed model (Model_saccade-LLH_) with the epochwise likelihood of each saccade type for each participant as an outcome variable. Within-subject predictors were Epoch (0-4) and saccade type (LDC, LDE, NLDC, NLDE).

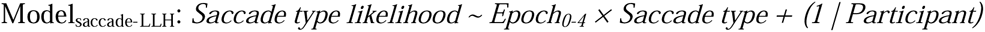

#### Iterative updating after different saccade types

To access the process of updating priors, we computed an iterative updating metric. For each trial, we first checked whether the participant had previously encountered the given triplet (e.g., 2-1-4). Trials featuring novel triplets were excluded. If the triplet had appeared earlier, we compared the participant’s predicted stimuli in the current and previous occurrences of that specific triplet (Figure 4). If the gaze was directed to the same location on both occasions, the trial was coded as ***no update***; if the predicted stimulus differed, it was coded as an ***update***. This yielded a binary variable (0 = no update, 1 = update) for each trial. For example, if a participant initially encountered the 2-1-4 triplet and looked toward the 4th circle before stimulus onset, and when the same 2-1-4 triplet occurred in the current trial, they looked toward the same location, this would be coded as no update. However, if they looked toward a different location (e.g., the 1st, 2nd, or 3rd circle) on the second occurrence, this would be coded as an update, indicating a change in their pre-stimulus expectation. We analyzed whether 1) the rate of updating changed throughout the task, 2) saccade type (LDC, LDE, NLDC, NLDE) moderated whether an update happened in the subsequent occurrence of the given triplet. To this end, we fit a generalized linear mixed model with a binomial distribution and logit link function (Model_saccade-update_). The binary outcome variable was trialwise Update (yes/no), and the within-subject predictors were Epoch (1-4) and previous saccade type (that is, the type the participant showed at the previous occurrence of the given triplet).

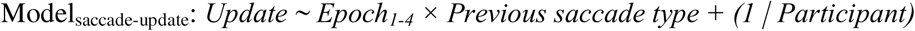

**Figure 4.**
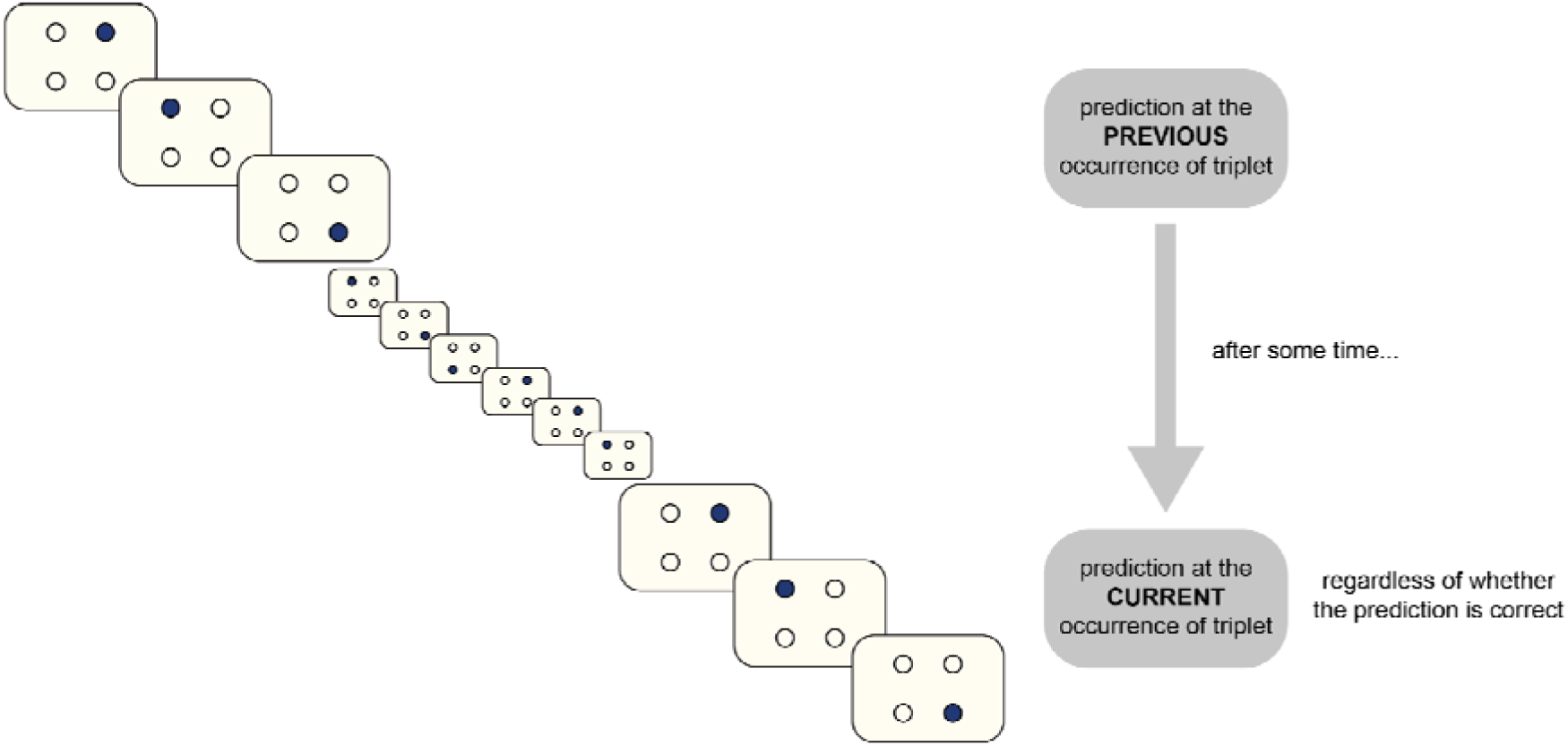
Schematic illustration of iterative updating. For each trial, we determined whether the presented triplet (e.g., 2-1-4) had occurred previously. Trials with novel triplets were excluded. For repeated triplets, the participant’s saccade at the current occurrence was compared to their saccade at the preceding occurrence of the same triplet. The schematic depicts this comparison between the previous and current occurrence of an identical triplet, irrespective of whether the earlier prediction was correct.

#### Likelihood of update types

To better understand what the updates reflect, we further categorized trials based on whether the presence or absence of updates presumably reflected learning. In trials where ***no update*** occurred – that is, the participant predicted the same stimulus as in the last occurrence of the triplet – we distinguished between ***LD same*** trials and ***NLD same*** trials. LD same trials were defined as instances where the participant exhibited a LD saccade in both occurrences, suggesting consistency with the statistical structure. NLD same trials, in contrast, were cases where the same NLD saccade was repeated, indicating persistence in an inaccurate or uninformative response.

In trials where an ***update*** occurred – that is, the participant changed their predicted stimulus – we further categorized the change based on the learning-dependence of the saccade in the previous and current occurrence of the given triplet. If a participant shifted from an LD to an NLD direction, we labeled the trial as an ***LD-to-NLD update***, reflecting a potential disruption in the internalized statistical representation. If the update went from an NLD to an LD direction, we categorized it as an ***NLD-to-LD update***, which we interpret as a possible correction toward the statistical structure. Finally, when both the previous and current saccade type was NLD, but the actual stimulus locations differed from each other, we labeled the trial as an ***NLD-to-other-NLD update***, which may reflect noise or exploratory behavior.

For instance, if the participant encountered the triplet 2-1-4 (which is high-probability in our above example sequence), and moved their eyes toward the 4th circle (LD saccade, as it corresponds to the high-probability triplet) both when it occurred previously and currently, we categorized the current one as an ***LD same*** trial. If they gazed toward the 3rd circle (a low-probability, NLD saccade) both times, we labeled it as an ***NLD same*** trial. If their saccade direction changed from the 4th to the 3rd circle between occurrences of the triplet 2-1-4, it was classified as an ***LD-to-NLD update***, whereas a shift from the 3rd to the 4th circle was classified as an ***NLD-to-LD update***. Lastly, if they first looked toward the 3rd circle and later toward the 1st or 2nd (both NLD saccades), we labeled the trial as an ***NLD-to-other-NLD update***.

Each of these categories provides insight into the participant’s learning process: LD same trials indicate the retention of correct statistical knowledge, NLD same trials reflect consistency in an incorrect representation, NLD-to-LD updates suggest the correction toward the statistical structure, LD-to-NLD updates point to a possible loss of learned information, and NLD-to-other-NLD updates likely reflect uncertainty or exploration.

To gain results that reflect the baseline probabilities, we divided the ratio of these update types in an epoch by their chance-level occurrence probabilities – similarly to the saccade type likelihoods, one reflects the chance level in this measurement. The baseline probability of these types is as follows: LD same: 0.0625, NLD same: 0.1875, LD-to-NLD: 0.1875, NLD-to-LD: 0.1875, NLD-to-other-NLD: 0.375, see Supplementary Table S3 for details.

To analyze the likelihoods of update types and how they changed throughout the task, we fit a linear mixed model (Model_update-LLH_) with the epochwise likelihood of each update type for each participant as an outcome variable. Within-subject predictors were Epoch (1-4) and update type (LD same, NLD same, LD-to-NLD update, NLD-to-LD update, NLD-to-other-NLD update).

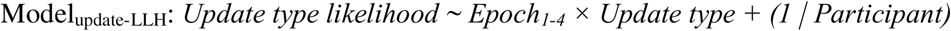

## Results

### 1. Do oRTs reflect statistical learning?

Here, we analyzed the trajectory of oculomotor reaction times (oRTs) in the task. oRTs refer to the latency from the appearance of a stimulus to the fixation on that stimulus. The difference in oRT between high- and low-probability triplets reflects statistical learning.

The linear mixed model testing the progress of learning in terms of oRTs (Model_SL-oRT_) reported a significant main effect of Epoch (*F*(4, 155.06) = 3.89, *p* = .005), which indicates a gradual increase of oRTs regardless of Triplet type (Epoch 0: 313 ms, 95% CI [310, 316]; Epoch 1: 315 ms, 95% CI [311, 318]; Epoch 2: 318 ms, 95% CI [313, 323]; Epoch 3: 318 ms, 95% CI [313, 322]; Epoch 4: 319 ms, 95% CI [315, 324]) (Figure 5). Pairwise contrasts between epochs are reported in Supplementary Table S4. The main effect of Triplet type was also significant (*F*(1, 762) = 120.07, *p* < .001), showing a generally slower response to low-compared to high-probability triplets across the task (Low-probability: 319 ms, 95% CI [315, 323]; High-probability: 314 ms, 95% CI [310, 318]; contrast *p_cor_ <* .001) (Figure 5). The significant Epoch × Triplet type interaction indicated a gradual change in the difference between low- and high-probability triplets (*F*(4, 762) = 9.54, *p* < .001), pairwise post-hoc comparisons revealing a progressive improvement in statistical learning (low-/high-probability differences: Epoch 0: 0.29 ms, *SE* = 1.05, *p_cor_* = .781; Epoch 1: 3.92 ms, *SE* = 1.05, *p_cor_* < .001; Epoch 2: 5.63 ms, *SE* = 1.05 *p_cor_* < .001; Epoch 3: 7.11 ms, *SE* = 1.05, *p_cor_* < .001; Epoch 4: 8.71 ms, *SE* = 1.05, *p_cor_* < .001) (Figure 5). The full list of fixed and random effect parameters is reported in Supplementary Table S5.

**Figure 5.**
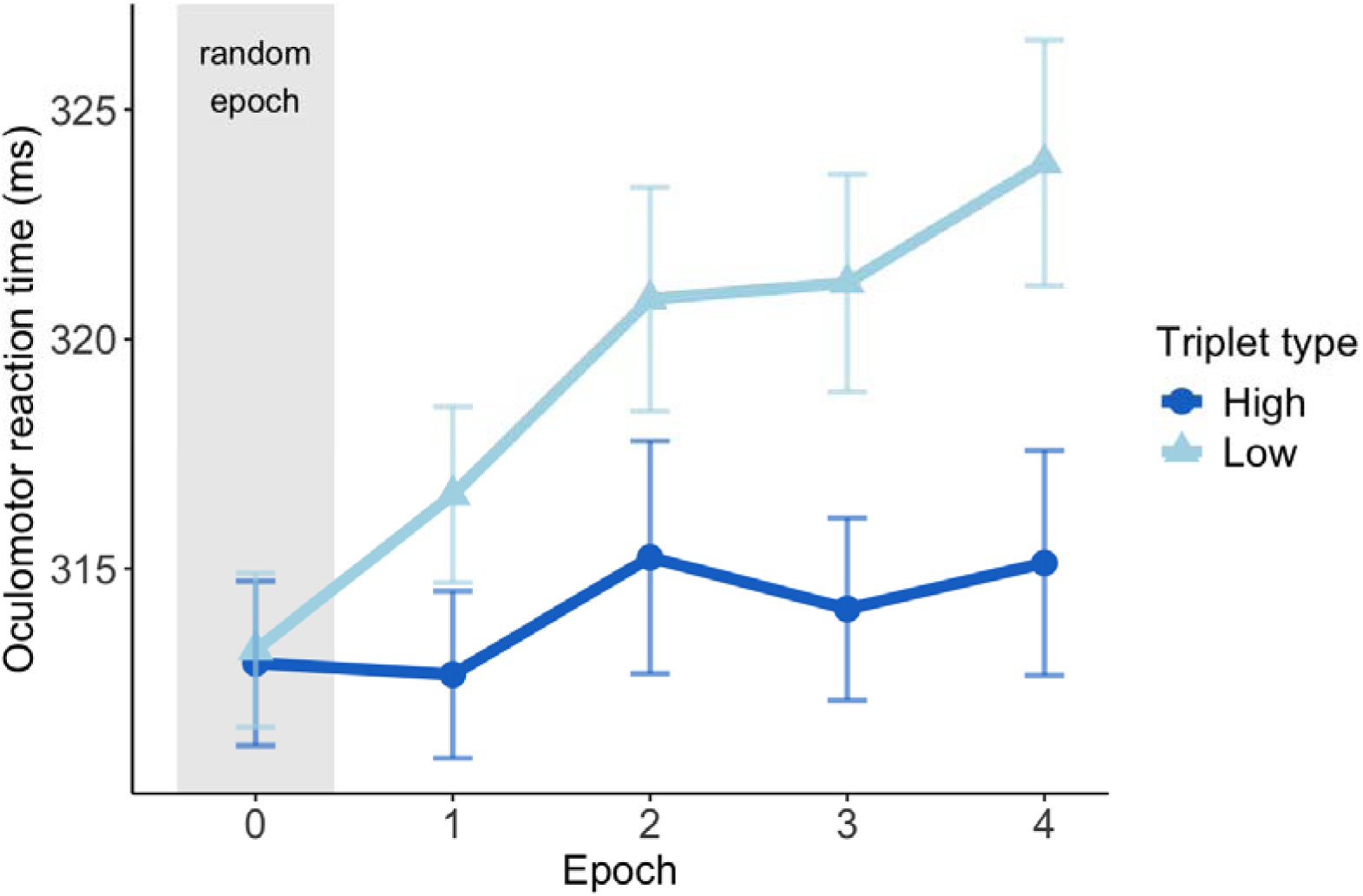
Statistical learning in the task. The x-axis represents epochs of the ASRT task (the progression of time). The y-axis represents performance in terms of oRTs (ms). The lines and markers show the trajectory of oRTs as a function of high-probability (dark blue line with circle markers) and low-probability (light blue line with triangle markers) triplets. The difference between high- and low-probability triplets represents statistical learning. Note that stimuli were presented randomly in Epoch 0 (marked as grey), serving as familiarization before the start of the actual task. Error bars show 1 SEM.

### 2. Do learning-dependent anticipation ratios reflect statistical learning?

Here, we analyzed the trajectory of learning-dependent anticipation ratios (LDARs) in the task. LDAR refers to the proportion of first saccades during the response-stimulus interval (RSI) directed toward the high-probability upcoming stimulus, relative to all first saccades of the trials (both toward high- and low-probability stimuli). Here, we did not further categorize these saccades as correct or erroneous. We used LDARs as an index of statistical learning, showing the extent to which first saccades are guided by the learned regularities of the task.

The linear mixed model analyzing the progress of learning in terms of LDARs (Model_SL-LDAR_) reported a significant main effect of Epoch (*F*(4, 508) = 48.73, *p* < .001), which reveals an increase in LDARs throughout the task (Epoch 0: 0.251, 95% CI = [0.243, 0.260]; Epoch 1: 0.290, 95% CI = [0.282, 0.299]; Epoch 2: 0.301, 95% CI = [0.293, 0.310]; Epoch 3: 0.305, 95% CI = [0.297, 0.314], Epoch 4: 0.314, 95% CI = [0.305, 0.322] (Figure 6). Pairwise contrasts between epochs are reported in Supplementary Table S6. The full list of fixed and random effect parameters is reported in Supplementary Table S7.

**Figure 6.**
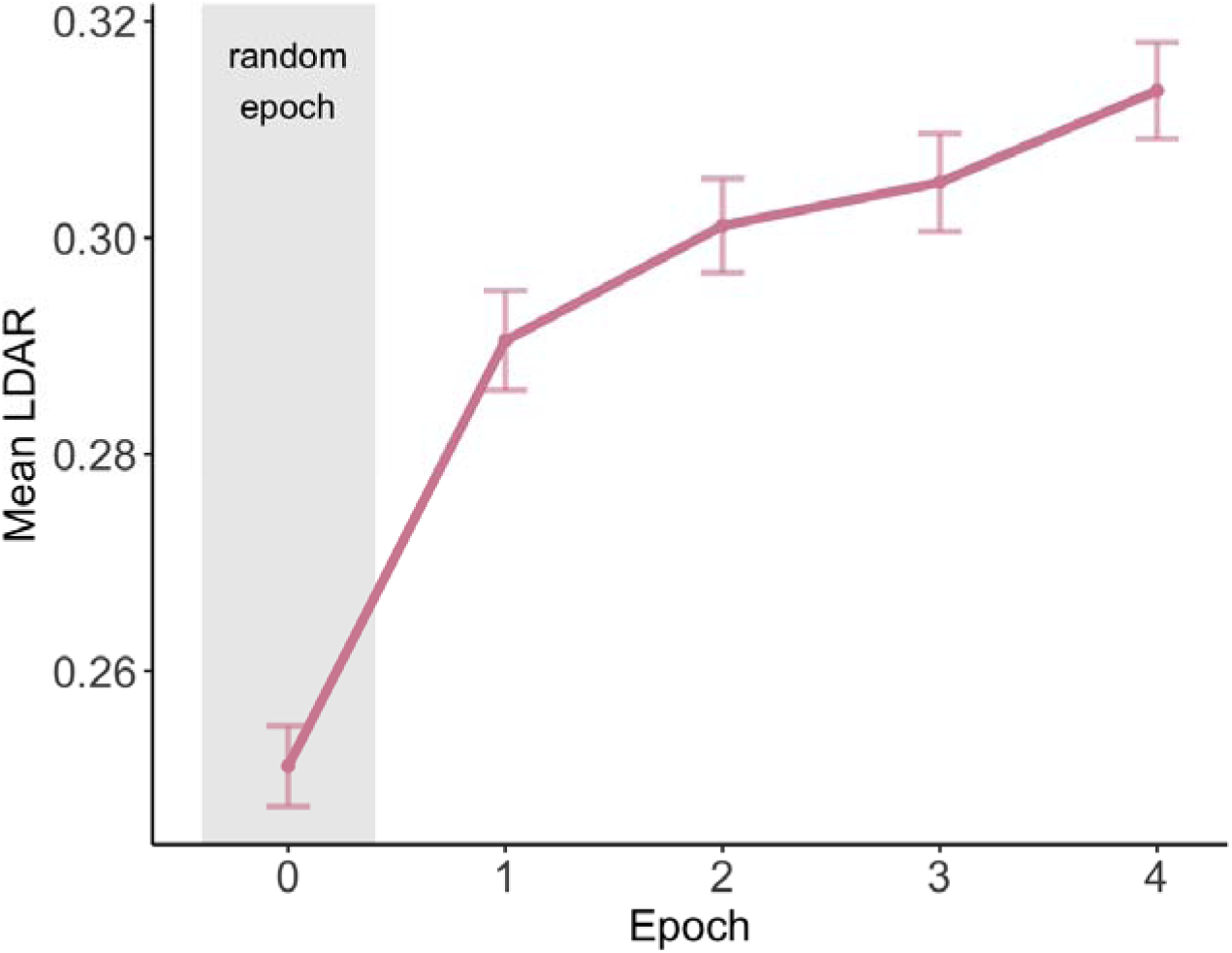
The ratio of learning-dependent anticipations in the task. The x-axis represents epochs of the ASRT task (the progression of time). The y-axis represents epochwise mean LDAR, the proportion of learning-dependent first saccades relative to all gaze shifts (that is, both toward high- and low-probability upcoming stimuli). The mauve line shows the trajectory of LDARs throughout the task. Note that stimuli were presented randomly in Epoch 0 (marked as grey), serving as familiarization before the start of the actual task. Error bars show 1 SEM.

### 3. Are learning-dependent errors more likely than not-learning-dependent errors?

Here, we analyzed the trajectory of the likelihood of saccade types in the task. We recorded the first saccade position after the last stimulus disappeared, categorized according to whether the participant’s gaze was directed toward the AOI of a high- or low-probability triplet, and whether this saccade matched the actual location of the upcoming stimulus. Saccades capture trial-by-trial predictive orienting behavior, reflecting whether expectations align with the statistical structure of the task.

The linear mixed model testing the likelihood of saccade types and their changes over time (Model_saccade-LLH_) showed a significant main effect of Epoch (*F*(4, 2413) = 13.124, *p* < .001), which indicates that the overall gradual change in likelihoods reached significance from Epoch 0 to 1, 2, 3 and 4. The model reported a significant main effect of Saccade type (*F*(3, 2413) = 263.762, *p* < .001), suggesting an overall difference in the likelihood of saccade types (Figure 7). LDEs were the most likely saccade type (1.239, 95% CI = [1.221, 1.256]), LDCs were the second most likely (1.124, 95% CI = [1.106, 1.141]), followed by NLDCs (0.948, 95% CI = [0.931, 0.966]), and, finally, NLDEs (0.944, 95% CI = [0.927, 0.962]; all pairwise contrasts between saccade types *p_cor_* < .001, except NLDC and NLDE *p_cor_*= .999). This difference in likelihoods was also modulated by Epoch, as indicated by a significant Epoch × Saccade type interaction (*F*(12, 2413) = 17.689, *p* < .001) (Figure 7). Based on model estimates and contrasts, LDEs significantly increased from Epoch 0 to all other epochs, and from Epoch 1 to 4. LDCs also increased from Epoch 0 to all other epochs, and from Epoch 1 to 4. NLDEs decreased from Epoch 0 to 4. NLDCs did not show a significant change across time. All post hoc model estimates and pairwise contrasts (by epoch and by saccade type, respectively) are reported in Supplementary Tables S8, S9, and S10, respectively, and the full list of fixed and random effect parameters is reported in Supplementary Table S11.

**Figure 7.**
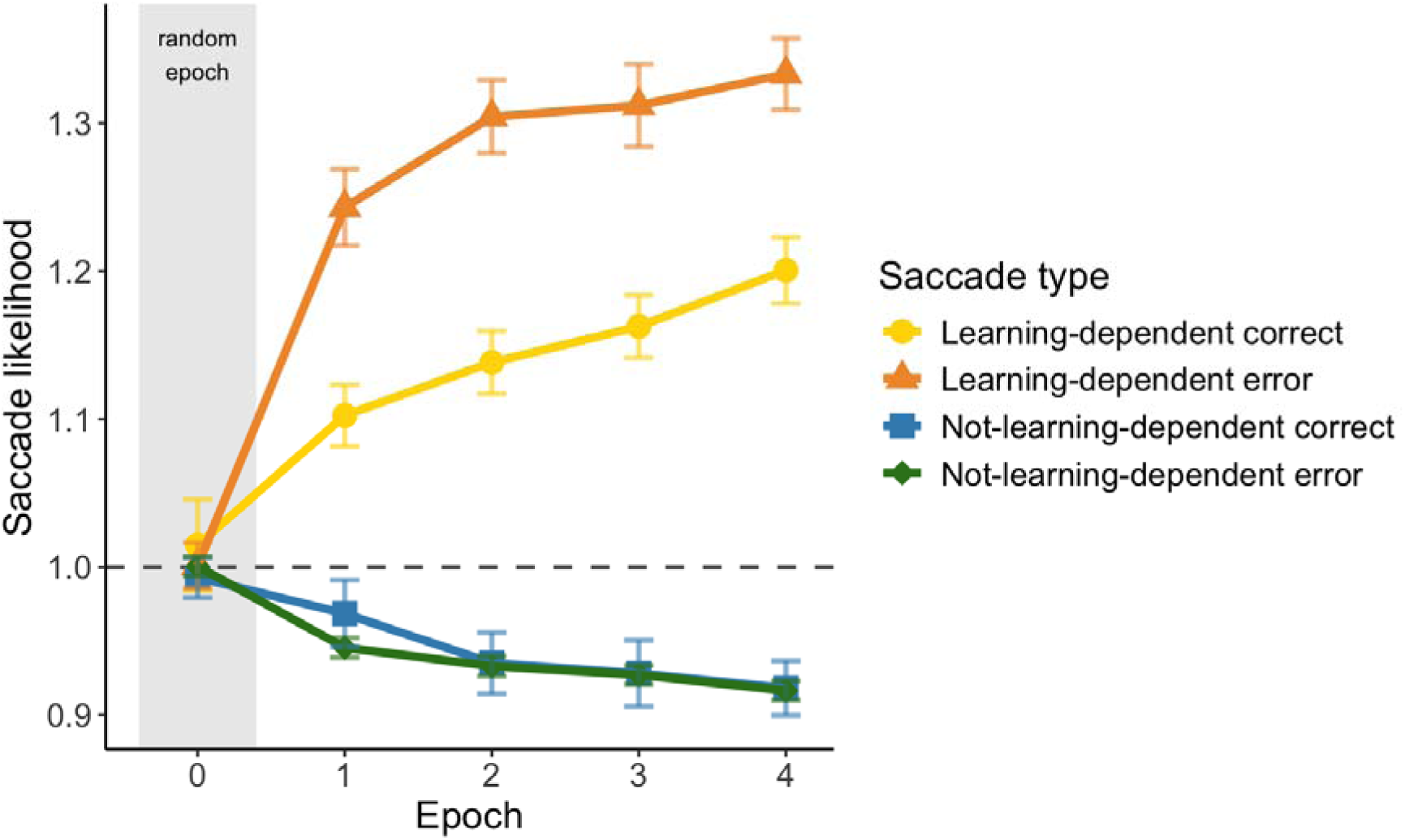
Likelihood of saccade types in the task. The x-axis represents epochs of the ASRT task (the progression of time). The y-axis represents the likelihood of saccade types, that is, their epochwise proportions standardized against their chance-level probabilities. As a result, a likelihood of 1 reflects chance-level probability (marked by a dashed grey line). The yellow line and circle markers represent the trajectory of the likelihood of learning-dependent correct saccade in the task, the orange line and triangle markers represent that of learning-dependent errors, the blue line and square markers represent that of not-learning-dependent corrects, and the green line and diamond markers represent that of not-learning-dependent errors. Note that stimuli were presented randomly in Epoch 0 (marked as grey), serving as familiarization before the start of the actual task. Error bars show 1 SEM.

### 4. Do participants update not-learning-dependent saccade types more often than learning-dependent saccade types?

Here, we analyzed how different saccade types were updated in the task. Update is a trialwise metric indicating whether the participant changed their saccade type for a given triplet relative to the most recent previous occurrence of that same triplet. No update means that the saccade type on the current trial is the same as it was on the previous occurrence of that triplet. The presence of an update means that the saccade type on the current trial differs from the previous occurrence of that triplet. Updating captures the dynamics of predictive adjustment across repeated presentations of the same triplet, reflecting stability or change in expectations. The epochwise raw number and percentage of updated vs. non-updated trials are shown in Supplementary Table S12.

The generalized linear mixed model investigating the iterative updating rate of different saccade types and how they changed throughout the task (Model_saccade-update_) reported a nonsignificant main effect of Epoch (□ ^2^(3) = 3.81, *p* = .283). (Figure 8). The main effect of Previous saccade type was significant (□ ^2^(3) = 257.89, *p* < .001), indicating that saccade types differed in the ratio at which they were updated, irrespective of time (Figure 8). The two most updated saccade types throughout the task were NLDCs (0.640, 95% CI = [0.624, 0.655]) and NLDE (0.630, 95% CI = [0.617, 0.642]). They did not significantly differ from one another (pairwise contrast *p_cor_* = .352), but they differed significantly from all other saccade types (all pairwise contrasts *p_cor_* < .001). The next most updated saccade type were LDCs (0.582, 95% CI = [0.567, 0.596]), which did not significantly differ from LDEs, the least updated saccade type (0.568, 95% CI = [0.553, 0.584]; pairwise contrast *p_cor_* = .133). The Epoch × Previous saccade type interaction was not significant (□ ^2^(9) = 16.81, *p* = .052), indicating no difference in the temporal trend of update ratios between saccade types (Figure 8). All post hoc model estimates and pairwise contrasts are reported in Supplementary Tables S13 and S14, and the full list of fixed and random effect parameters is reported in Supplementary Table S15.

**Figure 8.**
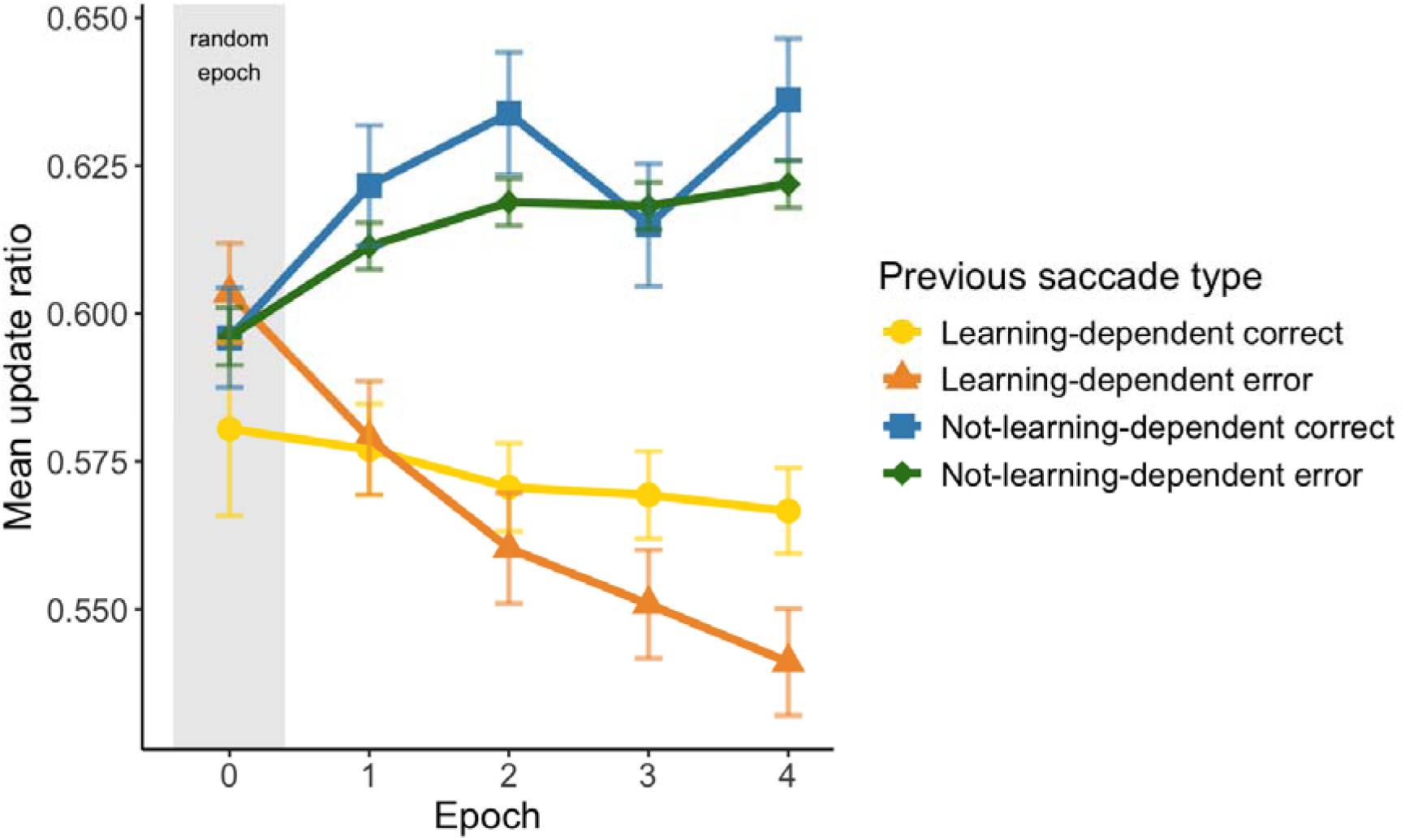
The ratio of the iterative updating of saccade types in the task. The x-axis represents epochs of the ASRT task (the progression of time). The y-axis represents epochwise mean update ratio, that is, the proportion of updated trials relative to all trials for each saccade type, separately. The yellow line and circle markers represent the trajectory of update ratios of learning-dependent correct saccades, the orange line and triangle markers represent that of learning-dependent errors, the blue line and square markers represent that of not-learning-dependent corrects, and the green line and diamond markers represent that of not-learning-dependent errors. Error bars show 1 SEM. Note that in this analysis, the practice (random) epoch is only presented in the visualization but could not be included in the analysis, as that would have altered the iterative measure derived from the analyzed data (since most triplets occurred the first time in this period).

Furthermore, we performed the same model with an additional fixed factor: the number of trials since the last occurrence of the given triplet to make sure that these results are not the artifact of (working) memory capacity. These analyses are reported in the Supplementary Table S16, S17, S18, S19, and Supplementary Figure S1.

### 5. Are updates that reflect learning more likely than updates that do not?

Here, we analyzed the likelihood of different update types in the task. Trials where an update occurs are further categorized based on the learning-dependence of the saccade type on the two occurrences of the given triplet. This yields five update types that suggest retention of correct statistical knowledge (LD same), persistence in an incorrect response (NLD same), potential loss or disruption of learned information (LD-to-NLD update), correction toward the statistical structure (NLD-to-LD update), or noise or exploratory behavior (NLD-to-other-NLD update). The epochwise raw number and percentage of trials in each update type are shown in Supplementary Table S20.

The linear mixed model analyzing the likelihood of update types and their changes over time (Model_update-LLH_) reported no significant main effect of Epoch (*F*(3, 2413) = 2.43, *p* = .064, indicating that there was no overall change in the likelihood of updates over time (Figure 9A). The main effect of Update type was significant (*F*(4, 2413) = 1011.01, *p* < .001), revealing an overall difference between the likelihood of different types of updates irrespective of time (Figure 9B). LD same trials were the most likely (2.026, 95% CI = [1.993, 2.059]), followed by NLD same trials (1.385, 95% CI = [1.353, 1.418]). Their likelihoods were significantly different from one another and all other update types (all pairwise contrasts involving LD same and/or NLD same *p_cor_* < .001). The next most likely update types were NLD-to-LD (0.934, 95% CI = [0.901, 0.967] and LD-to-NLD (0.905, 95% CI = [0.873, 0.938]), and their likelihoods significantly differed from all other update types (all pairwise contrasts *p_cor_* < .001) but not from one another (pairwise contrast *p_cor_* = .917). The least likely update type was NLD-to-other-NLD (0.717, 95% CI = [0.684, 0.750]), which differed significantly from all other update types (all pairwise contrasts *p_cor_*< .001). This difference in likelihoods was modulated by the interaction of Epoch and Update type (*F* (12, 2413) = 4.77, *p* < .001), indicating that Update type likelihoods followed a significantly different trend over time: the likelihood of LD same increased from Epoch 1 to 3 and 4 (Figure 9A). All post hoc model estimates and pairwise contrasts (by epoch and by saccade type, respectively) are reported in Supplementary Tables S21, S22, and S23, respectively, and the full list of fixed and random effect parameters is reported in Supplementary Table S24.

**Figure 9.**
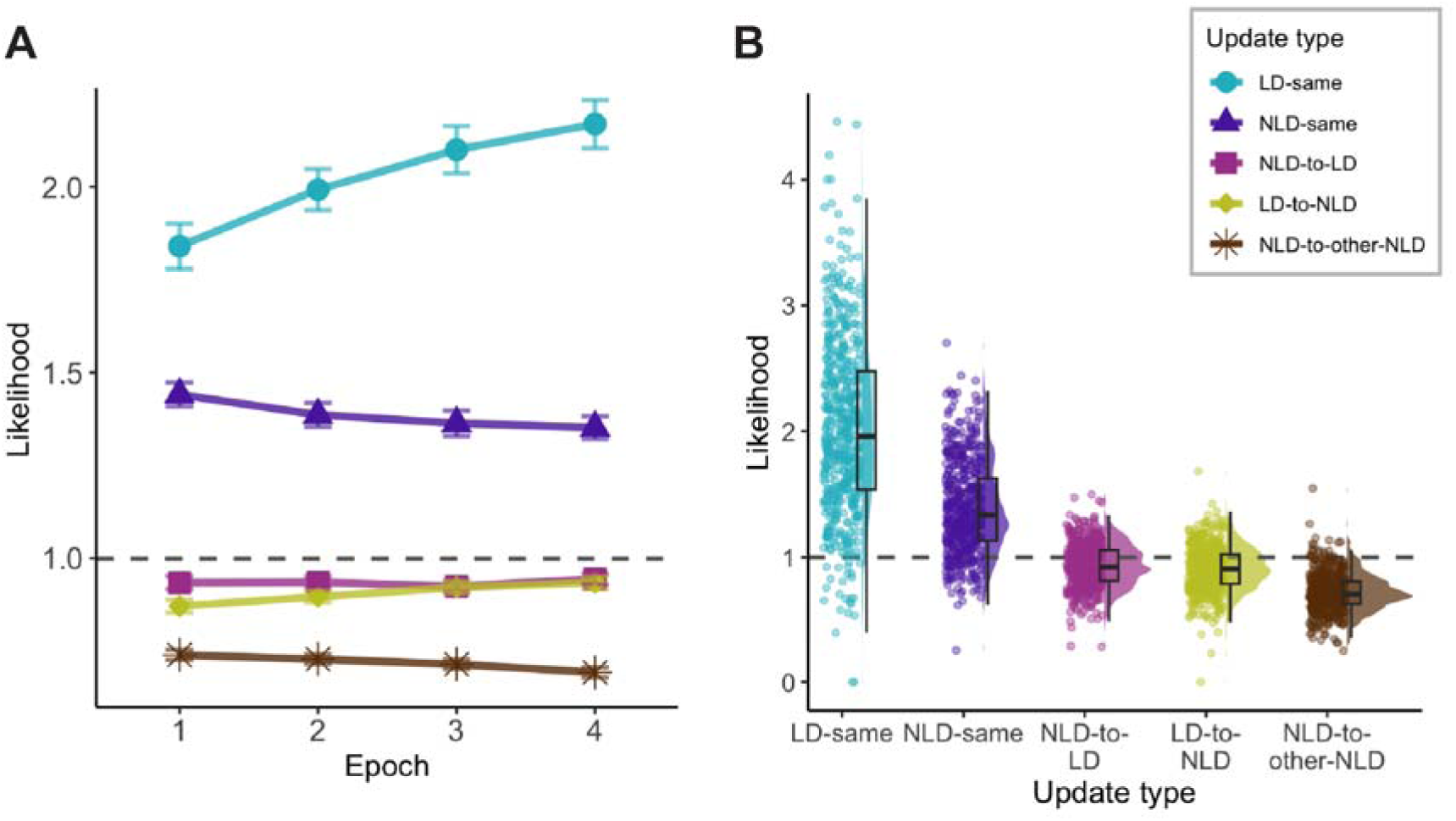
Likelihood of update types in the task. The y-axis represents the likelihood of update types, that is, their epochwise proportions standardized against their chance-level probabilities. As a result, a likelihood of 1 reflects chance-level probability (marked by a dashed grey line). Error bars show 1 SEM. **A)** Trajectories of update type likelihoods in the task. The x-axis represents epochs of the ASRT task (the progression of time). The turquoise line and round markers represent the trajectory of the likelihood of LD same occurrences in the task, the purple line and triangle markers represent that of NLD same occurrences, the pink line and square markers represent that of NLD-to-LD updates, the green line and diamond markers represent that of LD-to-NLD updates, and the brown line and star markers represent that of NLD-to-other-NLD updates. **B)** Distribution of likelihood values across update types. Each dot represents a single trial, with half-violins showing the density of the distribution and boxplots indicating the median and interquartile range. The x-axis represents the different update types.

## Discussion

To overcome the limitations of conventional offline and motor-based measures, here, we introduce a saccade-based analytical framework that provides a real-time window into the dynamics of prior expectations during statistical learning. We demonstrate this approach using a probabilistic sequence learning task, where we classify anticipatory saccades into two categories: learning-dependent anticipations, which align with the underlying statistical structure, and not-learning-dependent anticipations, which signal an inaccurate internal model. This distinction allows for the dissociation of prediction errors that arise from environmental stochasticity from those reflecting a lack of knowledge. Critically, we also introduce a metric to track the iterative updating of priors, identifying whether these changes reflect the consolidation of accurate predictions, the correction of errors, or exploratory behavior. Together, results using these measures suggest that statistical learning relies on conservative, repetition-based updating rather than uniform, strongly error-driven adjustments. The framework is paradigm-independent, offering a generalizable method for tracking the dynamic formation and refinement of predictive models in any task involving probabilistic stimulus streams.

By employing a highly sensitive, saccade-based analytical approach, we successfully captured the emergence and continuous growth of statistical learning from the earliest phases of exposure. Previous eye-tracking implementations of statistical learning paradigms are less sensitive: while detecting learning effects early, they often fail to capture the trajectory of learning later in the task. A recent study provided valuable insights into how learners actively form expectations about the probabilistic structure of their environment (Schwartz et al., 2026). They elegantly demonstrated how large-scale, block-by-block shifts in predictability can shape overall learning outcomes. While this macro-level analysis establishes that learners adapt to environmental noise over time, our findings dive into the real-time, micro-level dynamics behind this adaptation. Our framework allows us to track the gradual refinement of anticipatory behavior (i.e., learning-dependent anticipation ratios) over time. While this enhanced sensitivity is partially attributable to improved measurement precision, less data loss due to eye-tracking quality, and a larger sample size, it is fundamentally driven by our fine-grained analysis of pre-stimulus saccadic behavior. By isolating these subtle predictive eye movements, our approach provides a continuous, precise measure of underlying cognitive processes that standard spatial measures lack the resolution to detect.

Beyond the learning-dependentness of the saccade directions, we observed systematic patterns in erroneous predictions using our novel measurement. Learning-dependent errors (LDEs), or anticipations toward the high-probability but incorrect locations, were the most likely saccade type, and their likelihood significantly increased with time. In contrast, not-learning-dependent errors (NLDEs), i.e., erroneous saccades toward the low-probability and incorrect locations, remained below chance and showed a slight decrease throughout the task. This dissociation highlights that not all errors are equivalent, and that LDEs might reflect an engaged predictive system. In other words, the brain seems to differentiate between different error types, namely those that stem from noise and those that reflect inaccurate knowledge. LDEs – which reflect predictions congruent with the regularity but encounter noise – increase as the regularity is learned (reflected in oRTs and LDARs), despite the fact that they represent incorrect anticipations. The differentiation of error types highlights a central, unresolved issue of the statistical learning field: to what extent is statistical learning driven by errors, if at all? The fact that LDEs and NLDEs showed a largely different likelihood and opposing trajectory suggests that the brain uses different computations when one or the other occurs.

Furthermore, we found that participants updated their responses more frequently following NLDEs than LDEs. This suggests that NLDEs were assigned a higher learning rate (i.e., they were given a greater weight and elicited larger surprise) compared to LDEs. Importantly, this result remained stable when controlling for the number of trials elapsed since the last occurrence of the same event (see Supplementary Tables S16-S19 and Supplementary Figure S1), whereas the updating of learning-dependent correct trials seemed more distance-related. This suggests that the stronger impact of LDEs over NLDEs on updating cannot be explained simply by memory decay or working memory limitations. For such differential updating to occur, the brain may represent not merely the most probable outcome, but the full probability distribution of events – thereby allowing for calculated imprecision from the most probable option (i.e., LDEs). It suggests that learners encode expected uncertainty: they tolerate certain variability around the dominant pattern, rather than treating LDEs as signals that the regularity has changed (that is, unexpected uncertainty (Mathys et al., 2011) or volatility). Providing a behavioral measure to track the dissociation of these uncertainty types holds a practical and clinical importance, as alterations of this dissociation is often the marker of atypicalities and psychiatric conditions, such as in Autism Spectrum Disorder (Lawson et al., 2017; Palmer et al., 2017). Moreover, these findings suggest that computational models potentially assuming an unified learning rate for all prediction errors in probabilistic tasks may be prone to overlooking important asymmetries in updating. To obtain a deeper insight into iterative belief updating, future work could apply pupillometry, which offers quantifiable indices of surprise and precision-weighting.

Interestingly, within both learning-dependent and not-learning-dependent saccade types, correct and erroneous saccades did not differ in their likelihood of evoking updates upon the next occurrence of the same event (i.e., triplet). In other words, whether the anticipated stimulus actually appeared had little influence on subsequent adjustments. Instead, updating was more strongly determined by the structural alignment of the response with the task’s probabilistic regularities than by objective correctness. This finding suggests that statistical learning is not primarily governed by explicit confirmation or disconfirmation of predictions, but by the degree to which internal expectations are consistent with the underlying structure of the environment. Such a pattern is consistent with the notion that statistical learning operates in an unsupervised manner which may result in relative insensitivity to outcome feedback. More broadly, this result challenges the account that statistical learning is strongly driven by prediction errors. On the one hand, a Bayesian, error-driven approach is plausible under our results, if LDE and NLDE responses are attributed different weights (Fan et al., 2016; Friston et al., 2009; Olejarczuk et al., 2018). On the other hand, however, our results are consistent with the view that statistical learning is less driven by errors, but solely by the automatization of responses according to the task’s probability structure.

A more fine-grained analysis of the updating process revealed a strong repetition bias: participants tended to repeat their previous response to a given event. This repetition bias was more pronounced for learning-dependent responses (which also grew in likelihood as the task progressed), but it was also present for not-learning-dependent ones. Surprisingly, updates that corrected predictions inconsistent with the task’s probability structure (shifts from not-learning-dependent to learning-dependent responses) were similarly likely to update in the opposite direction. Thus, updating behavior did not primarily reflect a systematic correction of task-incongruent responses. Instead, once a response was produced, participants showed a strong tendency to maintain it across subsequent occurrences of the same event. This pattern points to a conservative updating style in statistical learning, plausible under two different interpretations. Interpreting it in a Bayesian framework, it means that rather than rapidly adjusting internal models after errors, learners appear to assign relatively low weight to individual prediction errors and instead stabilize previously formed expectations. Such a strategy may be adaptive in stable, non-volatile environments like the present task, where the underlying regularities remain constant over time (Simoens et al., 2024). Another plausible explanation is, again, that statistical learning is not error-driven.

This conservative updating profile can also be understood within reinforcement learning frameworks. From this perspective, anticipatory saccades and their subsequent adjustments reflect the balance between exploration and exploitation during learning (K. Friston, 2010; Mathys et al., 2011; Török et al., 2022). Not-learning-dependent updates may reflect exploration, where the system tests alternative predictions and helps the internal model to converge to the probabilistic structure (Divjak & Milin, 2020; Frank et al., 2021; Pesthy et al., 2025), while learning-dependent updates represent exploitation, using already established regularities (Howard & Howard, 1997; Kóbor et al., 2017; Németh et al., 2010; Song et al., 2007; Takács et al., 2018; Tóth-Fáber et al., 2023). Perseverative responses are not uncommon in the reinforcement learning literature, where similar patterns are often attributed to a choice-confirmation bias: learners tend to underweight belief-inconsistent evidence and preferentially integrate information that confirms their current expectations (Talluri et al., 2018). Although such processes are typically studied in reward-based paradigms, our findings suggest that similar reinforcement-like dynamics may operate in statistical learning even in the absence of explicit feedback. In this sense, the repetition bias observed here may reflect a general principle of adaptive learning systems: stabilizing predictions once they become sufficiently reliable, rather than continuously readjusting them after every error.

An alternative account emphasizes the habit-like nature of statistical learning. Repeating previously executed responses, even when they correspond to low-probability outcomes, is consistent with model-free mechanisms that rely on cached response tendencies rather than flexible, model-based representations (Daw et al., 2005; Poldrack et al., 2001). From this perspective, learners may partially encode the probabilistic structure of their own response patterns, rather than exclusively represent the statistical structure of the stimuli. In this sense, statistical learning may be driven not (or not only) by error-based updating, but also by the gradual response alignment with the regularity by a more Hebbian learning style. Presently, there is no consensus about how much errors matter in statistical learning – some studies support its Hebbian nature (Beesley & Shanks, 2012; Endress & Johnson, 2023; Tovar & Westermann, 2023; Vékony et al., 2022) while others found that it has a more error-driven nature (Fan et al., 2016; Leshinskaya & Thompson-Schill, 2018; Nazlı et al., 2024; Olejarczuk et al., 2018). Future studies should aim to systematically clarify this question, and our measurements potentially provide a tool for it.

From a Bayesian and predictive coding perspective, anticipatory eye movements are consistent with the notion of active inference, whereby the brain continuously generates predictions about upcoming events and minimizes prediction error by updating internal models (Friston, 2010; Friston et al., 2009). Within this framework, prediction errors are not weighted uniformly (Hohwy, 2020). Some arise from the probabilistic structure of the environment – reflecting expected uncertainty (Mathys et al., 2011) – whereas others indicate a mismatch between the internal model and the external world. The latter type of error is assigned greater precision and therefore exerts a stronger influence on model updating. This differential weighting may account for the observed differences between NLDEs and LDEs in our study. Importantly, participants showed a tendency to repeat their previous responses; in Bayesian terms, this suggests a stronger reliance on prior beliefs. In a stable, non-volatile environment such as ours, this strategy is adaptive, as previously acquired regularities are unlikely to change over time (Simoens et al., 2024). Thus, our results suggest that the Bayesian framework can help us understand the underlying processes of statistical learning (Éltető et al., 2022; Székely & Orbán, 2024; Török et al., 2022).

Some limitations of our study should be noted. First, our classification of anticipations into learning-dependent and not-learning-dependent categories assumes a single “correct” model of the task, i.e., the triplet-based second-order dependencies built into the ASRT. Although robust evidence indicates a group-level dissociation in performance on high-probability triplets in the ASRT task (Howard & Howard, 1997; Kóbor et al., 2017; Németh et al., 2010; Song et al., 2007; Takács et al., 2018; Tóth-Fáber et al., 2023), learners may not be building identical internal models, especially in the early stages of practice. Some participants may apply Markovian learning (Török et al., 2022), while others may form larger chunks of stimuli rather than triplets (Éltető et al., 2022; Szegedi-Hallgató et al., 2019). In such cases, an update we classified as not-learning-dependent relative to the model defined by the task could, in fact, be consistent with the participant’s own generative model. This means that our metric may underestimate the extent to which updates reflect rational model refinement from the learner’s perspective. Our measurement of model updating could also be biased by the fact that participants may be inclined to employ a conscious strategy of leaving their gaze at the previous stimulus position rather than reorienting it, resulting in saccades being present only in 48.76% of the trials. This could produce eye position carryover effects that don’t reflect internal model refinement. This strategy, however, is unlikely to significantly bias our likelihood results, as performance in the random epoch remained at the chance level. Future studies should aim to motivate participants to move their eyes, for example by using a stimulus stream that does not contain any repetitions, making leaving the gaze at the previous stimulus an ineffective strategy. Notably, we deliberately avoided the use of a central fixation cross between trials. Because the statistical regularity in the ASRT paradigm is built on the continuous sequential order of the stimuli, inserting a fixation cross would act as an additional sequence element, thereby transforming the transitional probabilities from second-order to third-order and fundamentally altering the learning mechanism. To further understand the mechanisms underlying statistical learning, our study points toward promising directions for future research. Future work could address the limitation of defining anticipations based on an externally defined “ground truth” model by combining saccade-based measures with computational modeling. Hierarchical Bayesian or active inference models (e.g., Mathys et al., 2011), for example, could be used to infer each participant’s latent hypothesis space, thereby allowing updates to be evaluated relative to the learner’s own evolving model rather than an a priori model defined by the task. Such an approach could strengthen the link between saccade dynamics and predictive coding accounts.

Our results provide two complementary contributions to the field of statistical learning. First, we developed a paradigm-independent methodological framework to capture the fine-grained dynamics of probabilistic learning, with a particular focus on the formation and updating of expectations. By isolating saccade-based markers of prediction, this approach offers a mechanistic account of learning processes that traditional motor-based measures overlook. Second, applying these new metrics allowed us a deeper insight into statistical learning. We show that learners differentiate between errors arising from probabilistic noise and those reflecting insufficient knowledge of the underlying structure. At the same time, within a given probability level, correct and erroneous predictions were equally likely to trigger subsequent updating, which is consistent with a less error-driven learning process that heavily relies on habitual repetition of previous responses.

## Data availability

All data are available at: https://osf.io/a4xhn/

## Code availability

The task script used in this study is available at: https://github.com/tzolnai/Child_ASRT_eye_tracking

All codes for data preprocessing, analysis, and visualization are available at: https://osf.io/a4xhn/

## References

1. Aslin, R. N. (2017). Statistical learning: A powerful mechanism that operates by mere exposure. Wiley Interdisciplinary Reviews: Cognitive Science, 8(1–2), e1373. 10.1002/wcs.1373

2. Bastos, A. M., Usrey, W. M., Adams, R. A., Mangun, G. R., Fries, P., & Friston, K. J. (2012). Canonical Microcircuits for Predictive Coding. Neuron, 76(4), 695–711. 10.1016/j.neuron.2012.10.038

3. Beesley, T., & Shanks, D. R. (2012). Investigating cue competition in contextual cuing of visual search. *Journal of Experimental Psychology: Learning*, Memory, and Cognition, 38(3), 709–725. 10.1037/a0024885

4. Chen, S., Zhang, X., Li, X., Jensen, O., Theeuwes, J., & Wang, B. (2025). Statistical learning drives anticipatory micro-saccades toward suppressed distractor locations (p. 2025.05.11.652318). bioRxiv. 10.1101/2025.05.11.652318

5. Clark, A. (2013). Whatever next? Predictive brains, situated agents, and the future of cognitive science. Behavioral and Brain Sciences, 36(3), 181–204. 10.1017/S0140525X12000477

6. Clark, A. (2016). Surfing Uncertainty: Prediction, Action, and the Embodied Mind. Oxford University Press. 10.1093/acprof:oso/9780190217013.001.0001

7. Compostella, A., Tagliani, M., Vender, M., & Delfitto, D. (2025). On the interaction between implicit statistical learning and the alternation advantage: Evidence from manual and oculomotor serial reaction time tasks. PLOS ONE, 20(2), e0318638. 10.1371/journal.pone.0318638

8. Daw, N. D., Niv, Y., & Dayan, P. (2005). Uncertainty-based competition between prefrontal and dorsolateral striatal systems for behavioral control. Nature Neuroscience, 8(12), 1704–1711. 10.1038/nn1560

9. Divjak, D., & Milin, P. (2020). Exploring and Exploiting Uncertainty: Statistical Learning Ability Affects How We Learn to Process Language Along Multiple Dimensions of Experience. Cognitive Science, 44(5), e12835. 10.1111/cogs.12835

10. Dodge, R. (1903). Five types of eye movement in the horizontal meridian plane of the field of regard. American Journal of Physiology-Legacy Content, 8(4), 307–329. 10.1152/ajplegacy.1903.8.4.307

11. Éltető, N., Nemeth, D., Janacsek, K., & Dayan, P. (2022). Tracking human skill learning with a hierarchical Bayesian sequence model. PLOS Computational Biology, 18(11), e1009866. 10.1371/journal.pcbi.1009866

12. Endress, A. D., & Johnson, S. P. (2023). Hebbian, correlational learning provides a memory-less mechanism for Statistical Learning irrespective of implementational choices: Reply to Tovar and Westermann (2022). Cognition, 230, 105290. 10.1016/j.cognition.2022.105290

13. Fan, J. E., Turk-Browne, N. B., & Taylor, J. A. (2016). Error-driven learning in statistical summary perception. Journal of Experimental Psychology. Human Perception and Performance, 42(2), 266–280. 10.1037/xhp0000132

14. Farkas, B. C., Krajcsi, A., Janacsek, K., & Nemeth, D. (2024). The complexity of measuring reliability in learning tasks: An illustration using the Alternating Serial Reaction Time Task. Behavior Research Methods, 56(1), 301–317. 10.3758/s13428-022-02038-5

15. Fox, J., & Weisberg, S. (2019). An R Companion to Applied Regression (Third). Sage. https://socialsciences.mcmaster.ca/jfox/Books/Companion/

16. Frank, S. M., Bründl, S., Frank, U. I., Sasaki, Y., Greenlee, M. W., & Watanabe, T. (2021). Fundamental Differences in Visual Perceptual Learning between Children and Adults. Current Biology, 31(2), 427–432.e5. 10.1016/j.cub.2020.10.047

17. Friston, K. (2005). A theory of cortical responses. Philosophical Transactions of the Royal Society B: Biological Sciences, 360(1456), 815–836. 10.1098/rstb.2005.1622

18. Friston, K. (2010). The free-energy principle: A unified brain theory? Nature Reviews Neuroscience, 11(2), 127–138. 10.1038/nrn2787

19. Friston, K. J., Daunizeau, J., & Kiebel, S. J. (2009). Reinforcement Learning or Active Inference? PLoS ONE, 4(7), e6421. 10.1371/journal.pone.0006421

20. Frost, R., Armstrong, B. C., & Christiansen, M. H. (2019). Statistical learning research: A critical review and possible new directions. Psychological Bulletin, 145(12), 1128–1153. 10.1037/bul0000210

21. Gregory, R. L. (1980). Perceptions as hypotheses. *Philosophical Transactions of the Royal Society of London. B*, Biological Sciences, 290(1038), 181–197. 10.1098/rstb.1980.0090

22. Harris, C. R., Millman, K. J., van der Walt, S. J., Gommers, R., Virtanen, P., Cournapeau, D., Wieser, E., Taylor, J., Berg, S., Smith, N. J., Kern, R., Picus, M., Hoyer, S., van Kerkwijk, M. H., Brett, M., Haldane, A., del Río, J. F., Wiebe, M., Peterson, P., … Oliphant, T. E. (2020). Array programming with NumPy. Nature, 585(7825), 357–362. 10.1038/s41586-020-2649-2

23. Hessels, R. S., Benjamins, J. S., van Doorn, A. J., Koenderink, J. J., Holleman, G. A., & Hooge, I. T. C. (2020). Looking behavior and potential human interactions during locomotion. Journal of Vision, 20(10), 5. 10.1167/jov.20.10.5

24. Hohwy, J. (2013). The Predictive Mind. Oxford University Press. 10.1093/acprof:oso/9780199682737.001.0001

25. Hohwy, J. (2020). New directions in predictive processing. Mind & Language, 35(2), 209–223. 10.1111/mila.12281

26. Howard, J. H., & Howard, D. V. (1997). Age differences in implicit learning of higher order dependencies in serial patterns. Psychology and Aging, 12(4), 634–656. 10.1037/0882-7974.12.4.634

27. Jiang, Y. V., Won, B.-Y., & Swallow, K. M. (2014). First saccadic eye movement reveals persistent attentional guidance by implicit learning. Journal of Experimental Psychology: Human Perception and Performance, 40(3), 1161–1173. 10.1037/a0035961

28. Keller, G. B., & Mrsic-Flogel, T. D. (2018). Predictive Processing: A Canonical Cortical Computation. Neuron, 100(2), 424–435. 10.1016/j.neuron.2018.10.003

29. Kelly, M. P., Kriznik, N. M., Kinmonth, A. L., & Fletcher, P. C. (2018). The brain, self and society: A social-neuroscience model of predictive processing. Social Neuroscience, 14(3), 266–276. 10.1080/17470919.2018.1471003

30. Knief, U., & Forstmeier, W. (2021). Violating the normality assumption may be the lesser of two evils. Behavior Research Methods, 53(6), 2576–2590. 10.3758/s13428-021-01587-5

31. Kóbor, A., Janacsek, K., Takács, Á., & Nemeth, D. (2017). Statistical learning leads to persistent memory: Evidence for one-year consolidation. Scientific Reports, 7(1), 760. 10.1038/s41598-017-00807-3

32. Lawson, R. P., Mathys, C., & Rees, G. (2017). Adults with autism overestimate the volatility of the sensory environment. Nature Neuroscience, 20(9), 1293–1299. 10.1038/nn.4615

33. Lenth, R. V. (2024). emmeans: Estimated Marginal Means, aka Least-Squares Means [Computer software]. https://CRAN.R-project.org/package=emmeans

34. Leshinskaya, A., & Thompson-Schill, S. L. (2018). Inferences about Uniqueness in Statistical Learning. Proceedings of the Annual Meeting of the Cognitive Science Society, 40(0). https://escholarship.org/uc/item/2bh0f7w8

35. Lieberman, M. D. (2000). Intuition: A social cognitive neuroscience approach. Psychological Bulletin, 126(1), 109–137. 10.1037/0033-2909.126.1.109

36. Lüdecke, D. (2023). sjPlot: Data Visualization for Statistics in Social Science [Computer software]. https://CRAN.R-project.org/package=sjPlot

37. Lüdecke, D., Ben-Shachar, M., Patil, I., Waggoner, P., & Makowski, D. (2021). performance: An R Package for Assessment, Comparison and Testing of Statistical Models. Journal of Open Source Software, 6(60), 3139. 10.21105/joss.03139

38. Lukács, Á., Lukics, K. S., & Dobó, D. (2021). Online statistical learning in developmental language disorder. Frontiers in Human Neuroscience, 15, 715818. 10.3389/fnhum.2021.715818

39. Lum, J. A. G. (2020). Incidental learning of a visuo-motor sequence modulates saccadic amplitude: Evidence from the serial reaction time task. Journal of Experimental Psychology. Learning, Memory, and Cognition, 46(10), 1881–1891. 10.1037/xlm0000917

40. Mathys, C., Daunizeau, J., Friston, K. J., & Stephan, K. E. (2011b). A Bayesian Foundation for Individual Learning Under Uncertainty. Frontiers in Human Neuroscience, 5. 10.3389/fnhum.2011.00039

41. McKinney, W. (2010). Data Structures for Statistical Computing in Python. 56–61. 10.25080/Majora-92bf1922-00a

42. Miłkowski, M., & Litwin, P. (2022). Testable or bust: Theoretical lessons for predictive processing. Synthese, 200(6), 462. 10.1007/s11229-022-03891-9

43. Nakagawa, S., & Schielzeth, H. (2013). A general and simple method for obtaining R2 from generalized linear mixed-effects models. Methods in Ecology and Evolution, 4(2), 133–142. 10.1111/j.2041-210x.2012.00261.x

44. Nazlı, İ., Ferrari, A., Huber-Huber, C., & Lange, F. P. de. (2024). Forward and backward blocking in statistical learning. PLOS ONE, 19(8), e0306797. 10.1371/journal.pone.0306797

45. Németh, D., Janacsek, K., Balogh, V., Londe, Z., Mingesz, R., Fazekas, M., Jambori, S., Danyi, I., & Vetro, A. (2010). Learning in autism: Implicitly superb. PLoS ONE, 5(7), e11731. 10.1371/journal.pone.0011731

46. Németh, D., Janacsek, K., & Fiser, J. (2013). Age-dependent and coordinated shift in performance between implicit and explicit skill learning. Frontiers in Computational Neuroscience, 7. 10.3389/fncom.2013.00147

47. Németh, D., & Tóth-Fáber, E. (2026). Resolving Fundamental Debates in Statistical Learning: The Power of Process Dissociation. PsyArXiv. https://osf.io/uwa35_v1

48. Németh, D., Tóth-Fáber, E., Farkas, B., & Janacsek, K. (2026). Revisiting Age-Related Changes in Statistical Learning: The Importance of Longitudinal Evidence. Cognitive Science, 50(1), e70174. 10.1111/cogs.70174

49. Nissen, M. J., & Bullemer, P. (1987). Attentional requirements of learning: Evidence from performance measures. Cognitive Psychology, 19(1), 1–32. 10.1016/0010-0285(87)90002-8

50. Nyström, M., & Holmqvist, K. (2010). An adaptive algorithm for fixation, saccade, and glissade detection in eyetracking data. Behavior Research Methods, 42(1), 188–204. 10.3758/BRM.42.1.188

51. Olejarczuk, P., Kapatsinski, V., & Baayen, R. H. (2018). Distributional learning is error-driven: The role of surprise in the acquisition of phonetic categories. Linguistics Vanguard, 4(s2). 10.1515/lingvan-2017-0020

52. Palmer, C. J., Lawson, R. P., & Hohwy, J. (2017). Bayesian approaches to autism: Towards volatility, action, and behavior. Psychological Bulletin, 143(5), 521–542. 10.1037/bul0000097

53. Peirce, J., Gray, J. R., Simpson, S., MacAskill, M., Höchenberger, R., Sogo, H., Kastman, E., & Lindeløv, J. K. (2019). PsychoPy2: Experiments in behavior made easy. Behavior Research Methods, 51(1), 195–203. 10.3758/s13428-018-01193-y

54. Pesthy, O., Pesthy, Z. V., Vékony, T., Fabó, D., & Nemeth, D. (2025). Inhibiting the right dorsolateral prefrontal cortex selectively enhances unsupervised statistical learning (p. 2025.08.08.669288). bioRxiv. 10.1101/2025.08.08.669288

55. Poldrack, R. A., Clark, J., Paré-Blagoev, E. J., Shohamy, D., Creso Moyano, J., Myers, C., & Gluck, M. A. (2001). Interactive memory systems in the human brain. Nature, 414(6863), 546–550. 10.1038/35107080

56. Project ET Developers. (2021). *Project ET Zero* (Version 0.1.0) [Python]. https://github.com/tzolnai/project_ET_zero (Original work published 2020)

57. Python Software Foundation. (2024). The Python Language Reference (Version 3.13) [Computer software]. https://docs.python.org/3/reference/index.html

58. R Core Team. (2024). R: A language and environment for statistical computing [Computer software]. Vienna, Austria: R Foundation for Statistical Computing. https://www.R-project.org/

59. Rao, R. P. N., & Ballard, D. H. (1999). Predictive coding in the visual cortex: A functional interpretation of some extra-classical receptive-field effects. Nature Neuroscience, 2(1), 79–87. 10.1038/4580

60. Romano Bergstrom, J. C., Howard Jr., J. H., & Howard, D. V. (2012). Enhanced Implicit Sequence Learning in College-age Video Game Players and Musicians. Applied Cognitive Psychology, 26(1), 91–96. 10.1002/acp.1800

61. Satterthwaite, F. E. (1941). Synthesis of variance. Psychometrika, 6(5), 309–316. 10.1007/BF02288586

62. Schielzeth, H., Dingemanse, N. J., Nakagawa, S., Westneat, D. F., Allegue, H., Teplitsky, C., Réale, D., Dochtermann, N. A., Garamszegi, L. Z., & ArayaLJAjoy, Y. G. (2020). Robustness of linear mixedLJeffects models to violations of distributional assumptions. Methods in Ecology and Evolution, 11(9), 1141–1152. 10.1111/2041-210X.13434

63. Schlichting, M. L., Guarino, K. F., Schapiro, A. C., Turk-Browne, N. B., & Preston, A. R. (2017). Hippocampal Structure Predicts Statistical Learning and Associative Inference Abilities during Development. Journal of Cognitive Neuroscience, 29(1), 37–51. 10.1162/jocn_a_01028

64. Schwartz, N., Bogaerts, L., Loyfer, Y., Tal, A., Siegelman, N., & Frost, R. (2026). Statistical learning performance is impacted by a previous learning experience: A predictive eye-movement study. Cognition, 272, 106512. 10.1016/j.cognition.2026.106512

65. Šidák, Z. (1967). Rectangular Confidence Regions for the Means of Multivariate Normal Distributions. Journal of the American Statistical Association, 62(318), 626–633. 10.1080/01621459.1967.10482935

66. Siegelman, N., Bogaerts, L., Kronenfeld, O., & Frost, R. (2018). Re-defining “learning” in statistical learning: What does an online measure reveal about the assimilation of visual regularities? Cognitive Science, 42(Suppl 3), 692–727. 10.1111/cogs.12556

67. Simoens, J., Verguts, T., & Braem, S. (2024). Learning environment-specific learning rates. PLOS Computational Biology, 20(3), e1011978. 10.1371/journal.pcbi.1011978

68. Simor, P., Zavecz, Z., Horváth, K., Éltető, N., Török, C., Pesthy, O., Gombos, F., Janacsek, K., & Nemeth, D. (2019). Deconstructing Procedural Memory: Different Learning Trajectories and Consolidation of Sequence and Statistical Learning. Frontiers in Psychology, 9. 10.3389/fpsyg.2018.02708

69. Singmann, H., Bolker, B., Westfall, J., Aust, F., & Ben-Shachar, M. S. (2024). afex: Analysis of Factorial Experiments [Computer software]. https://CRAN.R-project.org/package=afex

70. Soetens, E., Melis, A., & Notebaert, W. (2004). Sequence learning and sequential effects. Psychological Research, 69(1–2), 124–137. 10.1007/s00426-003-0163-4

71. Song, S., Howard, J. H., & Howard, D. V. (2007). Sleep Does Not Benefit Probabilistic Motor Sequence Learning. The Journal of Neuroscience, 27(46), 12475. 10.1523/JNEUROSCI.2062-07.2007

72. Szegedi-Hallgató, E., Janacsek, K., & Nemeth, D. (2019). Different levels of statistical learning—Hidden potentials of sequence learning tasks. PLoS ONE, 14(9). 10.1371/journal.pone.0221966

73. Szegedi-Hallgató, E., Janacsek, K., Vékony, T., Tasi, L. A., Kerepes, L., Hompoth, E. A., Bálint, A., & Németh, D. (2017). Explicit instructions and consolidation promote rewiring of automatic behaviors in the human mind. Scientific Reports, 7(1), 4365. 10.1038/s41598-017-04500-3

74. Székely, A., & Orbán, G. (2024). Bayes beyond the predictive distribution. The Behavioral and Brain Sciences, 47, e166. 10.1017/S0140525X24000086

75. Takács, Á., Kóbor, A., Chezan, J., Éltető, N., Tárnok, Z., Nemeth, D., Ullman, M. T., & Janacsek, K. (2018). Is procedural memory enhanced in Tourette syndrome? Evidence from a sequence learning task. Cortex, 100, 84–94. 10.1016/j.cortex.2017.08.037

76. Takacs, A., Toth-Faber, E., Schubert, L., Tarnok, Z., Ghorbani, F., Trelenberg, M., Nemeth, D., Münchau, A., & Beste, C. (2024). Neural representations of statistical and rule-based predictions in Gilles de la Tourette syndrome. Human Brain Mapping, 45(8), e26719. 10.1002/hbm.26719

77. Tal, A., Bloch, A., Cohen-Dallal, H., Aviv, O., Schwizer Ashkenazi, S., Bar, M., & Vakil, E. (2021). Oculomotor anticipation reveals a multitude of learning processes underlying the serial reaction time task. Scientific Reports, 11(1), 6190. 10.1038/s41598-021-85842-x

78. Tal, A., & Vakil, E. (2020). How sequence learning unfolds: Insights from anticipatory eye movements. Cognition, 201, 104291. 10.1016/j.cognition.2020.104291

79. Talluri, B. C., Urai, A. E., Tsetsos, K., Usher, M., & Donner, T. H. (2018). Confirmation Bias through Selective Overweighting of Choice-Consistent Evidence. Current Biology, 28(19), 3128–3135.e8. 10.1016/j.cub.2018.07.052

80. team, T. pandas development. (2026). pandas-dev/pandas: Pandas (Version v3.0.1) [Computer software]. Zenodo. 10.5281/zenodo.18675244

81. Tobii AB. (2020a). Data quality reports for 3 Tobii eye trackers. https://www.tobii.com/resource-center/data-quality

82. Tobii AB. (2020b). Tobii Pro SDK for Python (Version 1.8.0) [Computer software]. https://pypi.org/project/tobii-research/

83. Tobii AB. (2024). Tobii Pro Eye Tracker Manager [Computer software]. https://developer.tobiipro.com/eyetrackermanager.html

84. Tobii AB. (2025). Tobii Pro Fusion eye tracker: Product description. https://www.tobii.com/products/eye-trackers/screen-based/tobii-pro-fusion

85. Török, B., Nagy, D. G., Kiss, M., Janacsek, K., Németh, D., & Orbán, G. (2022). Tracking the contribution of inductive bias to individualised internal models. PLOS Computational Biology, 18(6), e1010182. 10.1371/journal.pcbi.1010182

86. Tóth-Fáber, E., Németh, D., & Janacsek, K. (2023). Lifespan developmental invariance in memory consolidation: Evidence from procedural memory. PNAS Nexus, 2(3), pgad037. 10.1093/pnasnexus/pgad037

87. Tovar, Á. E., & Westermann, G. (2023). No need to forget, just keep the balance: Hebbian neural networks for statistical learning. Cognition, 230, 105176. 10.1016/j.cognition.2022.105176

88. Ullman, M. T. (2016). The Declarative/Procedural Model: A Neurobiological Model of Language Learning, Knowledge, and Use. In G. Hickok & S. L. Small (Eds.), Neurobiology of Language (pp. 953–968). Academic Press. 10.1016/B978-0-12-407794-2.00076-6

89. Vakil, E., Bloch, A., & Cohen, H. (2017). Anticipation Measures of Sequence Learning: Manual versus Oculomotor Versions of the Serial Reaction Time Task. Quarterly Journal of Experimental Psychology, 70(3), 579–589. 10.1080/17470218.2016.1172095

90. Vékony, T., Ambrus, G. G., Janacsek, K., & Nemeth, D. (2022). Cautious or causal? Key implicit sequence learning paradigms should not be overlooked when assessing the role of DLPFC (Commentary on Prutean et al.). Cortex, 148, 222–226. 10.1016/j.cortex.2021.10.001

91. Vékony, T., Pleche, C., Pesthy, O., Janacsek, K., & Nemeth, D. (2022). Speed and accuracy instructions affect two aspects of skill learning differently. Npj Science of Learning, 7(1), 27. 10.1038/s41539-022-00144-9

92. Vékony, T., Takács, Á., Pedraza, F., Haesebaert, F., Tillmann, B., Mihalecz, I., Phelipon, R., Beste, C., & Nemeth, D. (2023). Modality-specific and modality-independent neural representations work in concert in predictive processes during sequence learning. Cerebral Cortex, 33(12), 7783–7796. 10.1093/cercor/bhad079

93. Virtanen, P., Gommers, R., Oliphant, T. E., Haberland, M., Reddy, T., Cournapeau, D., Burovski, E., Peterson, P., Weckesser, W., Bright, J., van der Walt, S. J., Brett, M., Wilson, J., Millman, K. J., Mayorov, N., Nelson, A. R. J., Jones, E., Kern, R., Larson, E., … SciPy 1.0 Contributors. (2020). SciPy 1.0: Fundamental algorithms for scientific computing in Python. Nature Methods, 17(3), 261–272. 10.1038/s41592-019-0686-2

94. Westheimer, G. (1954). Mechanism of saccadic eye movements. A.M.A. Archives of Ophthalmology, 52, 710–723. 10.1001/archopht.1954.00920050716006

95. Wickham, H. (2016). ggplot2: Elegant Graphics for Data Analysis. Springer-Verlag New York. https://ggplot2.tidyverse.org

96. Williams, D. (2018). Predictive processing and the representation wars. Minds and Machines, 28(1), 141–172. 10.1007/s11023-017-9441-6

97. Yu, C., & Smith, L. B. (2011). What you learn is what you see: Using eye movements to study infant cross-situational word learning. Developmental Science, 14(2), 165–180. 10.1111/j.1467-7687.2010.00958.x

98. Yu, C., Zhong, Y., & Fricker, D. (2012). Selective Attention in Cross-Situational Statistical Learning: Evidence From Eye Tracking. Frontiers in Psychology, 3. 10.3389/fpsyg.2012.00148

99. Zolnai, T., Dávid, D. R., Pesthy, O., Nemeth, M., Kiss, M., Nagy, M., Nemeth, D., & Ergul, A. (2022). Measuring statistical learning by eye-tracking. Experimental Results, 3, e10. 10.1017/exp.2022.8

