## Supplementary Materials for "Capturing learning on the fly: an eye-tracking method to quantify prediction errors and updating the prior"

### Descriptive statistics of saccades

**Supplementary Table S1.** Descriptive statistics of the general metrics of saccade data. Saccades with latency longer than 500 ms (M_count/subject_ = 0.73 trials, SD = 1.18 trials) were dropped as they signal PsychoPy malfunction.

| **Metric** | **Mean** | **Standard Deviation** |
| --- | --- | --- |
| Velocity (°/s) | 140.98 | 75.32 |
| Amplitude (°) | 6.58 | 5.05 |
| Duration (ms) | 46.10 | 20.14 |
| Latency (ms) | 229.03 | 101.56 |

### Baseline probabilities of saccade types

We calculated the baseline (chance-level) probabilities of saccade types based on the chance-level probability of the saccade type for each location (P = 0.25 for each of the four circles), and the chance-level probability of all stimulus outcomes (for high-probability triplets, P = 0.635, and for each of the three low-probability triplet options, P = 0.125). The calculated probabilities are shown in Supplementary Table S2.

**Supplementary Table S2.** Baseline (chance-level) probabilities of each saccade type.

| **Saccade direction** | **Actual outcome** | **Correct** | **P of event** | **Label** |
| --- | --- | --- | --- | --- |
| High-prob (P = 0.25) | High-prob (P = 0.625) | Yes | 0.15625 | LDC |
|  | Low-prob/1 (P = 0.125) | No | 0.03125 | LDE |
|  | Low-prob/2 (P = 0.125) | No | 0.03125 | LDE |
|  | Low-prob/3 (P = 0.125) | No | 0.03125 | LDE |
| Low-prob/1 (P = 0.25) | High-prob (P = 0.625) | No | 0.15625 | NLDE |
|  | Low-prob/1 (P = 0.125) | Yes | 0.03125 | NLDC |
|  | Low-prob/2 (P = 0.125) | No | 0.03125 | NLDE |
|  | Low-prob/3 (P = 0.125) | No | 0.03125 | NLDE |
| Low-prob/2 (P = 0.25) | High-prob (P = 0.125) | No | 0.15625 | NLDE |
|  | Low-prob/1 (P = 0.125) | No | 0.03125 | NLDE |
|  | Low-prob/2 (P = 0.125) | Yes | 0.03125 | NLDC |
|  | Low-prob/3 (P = 0.125) | No | 0.03125 | NLDE |
| Low-prob/3 (P = 0.25) | High-prob (P = 0.125) | No | 0.15625 | NLDE |
|  | Low-prob/1 (P = 0.125) | No | 0.03125 | NLDE |
|  | Low-prob/2 (P = 0.125) | No | 0.03125 | NLDE |
|  | Low-prob/3 (P = 0.125) | Yes | 0.03125 | NLDC |
| Total LDC | | | 0.15625 |  |
| Total LDE | | | 0.09375 |  |
| Total NLDC | | | 0.09375 |  |
| Total NLDE | | | 0.65625 |  |
| Total | | | 1 |  |

*Note.* High-prob = the last element of a high-probability triplet, low-prob/1, 2, or 3: the last element of one of the possible low-probability triplets. Ps reflect the chance-level probability of a given outcome. LDC = learning-dependent correct, LDE = learning-dependent error, NLDC = not-learning-dependent correct, NLDE = not-learning-dependent error.

### Baseline probabilities of update types

The calculation of update type chance-level probabilities happened as follows. We calculated the joint probability of (a) a given saccade type to happen (see above in Table S1), (b) the same triplet to occur again (see Methods/Task and procedure for details), and (c) a specific saccade type to happen at the current occurrence. This resulted in the chance-level probabilities demonstrated in Table S3.

**Supplementary Table S3.** Baseline (chance-level) probabilities of each update type.

| **Update type** | P |
| --- | --- |
| LD same | 0.0625 |
| NLD same | 0.1875 |
| LD-to-NLD update | 0.1875 |
| NLD-to-LD update | 0.1875 |
| NLD-to-other-NLD update | 0.375 |

*Note.* P refers to the baseline (chance-level) probability of each event.

### Model_SL-oRT_

**Supplementary Table S4.** Contrasts by epoch between the estimates of oRT.

| **Contrast** | ***p*** |
| --- | --- |
| Epoch 0 - Epoch 1 | 0.603 |
| Epoch 0 - Epoch 2 | **0.046** |
| Epoch 0 - Epoch 3 | **0.036** |
| Epoch 0 - Epoch 4 | **0.006** |
| Epoch 1 - Epoch 2 | 0.201 |
| Epoch 1 - Epoch 3 | 0.081 |
| Epoch 1 - Epoch 4 | **0.008** |
| Epoch 2 - Epoch 3 | 1.000 |
| Epoch 2 - Epoch 4 | 0.996 |
| Epoch 3 - Epoch 4 | 0.723 |

**Supplementary Table S5.** Results of the linear mixed model predicting epochwise mean oRT.

| **Prediction of oRT** | | | | | |
| --- | --- | --- | --- | --- | --- |
| **Fixed Effects** | ***β estimates*** | ***95% CI*** | ***t*** | ***df*** | ***p*** |
| **(Intercept)** | 316.588 | 312.804 – 320.372 | 165.542 | 127.000 | **<0.001** |
| **Epoch [1]** | -3.509 | -5.515 – -1.503 | -3.461 | 127.951 | **0.001** |
| **Epoch [2]** | -1.938 | -3.148 – -0.728 | -3.162 | 162.609 | **0.002** |
| Epoch [3] | 1.471 | -0.773 – 3.715 | 1.297 | 127.003 | 0.197 |
| Epoch [4] | 1.084 | -0.496 – 2.664 | 1.358 | 127.458 | 0.177 |
| **Triplet type [high]** | -2.566 | -3.025 – -2.106 | -10.957 | 762.000 | **<0.001** |
| **Epoch [1] × Triplet type [high]** | 2.420 | 1.501 – 3.339 | 5.168 | 762.000 | **<0.001** |
| Epoch [2] × Triplet type [high] | 0.605 | -0.315 – 1.524 | 1.291 | 762.000 | 0.197 |
| Epoch [3] × Triplet type [high] | -0.247 | -1.166 – 0.672 | -0.528 | 762.000 | 0.598 |
| **Epoch [4] × Triplet type [high]** | -0.987 | -1.907 – -0.068 | -2.108 | 762.000 | **0.035** |
| **Random Effects** | | | | | |
| σ^2^ | 70.177 |  |  |  |  |
| τ_00_ _id_ | 461.130 |  |  |  |  |
| τ_11_ _id.epoch1_ | 103.525 |  |  |  |  |
| τ_11_ _id.epoch2_ | 19.999 |  |  |  |  |
| τ_11_ _id.epoch3_ | 136.551 |  |  |  |  |
| τ_11_ _id.epoch4_ | 53.519 |  |  |  |  |
| ρ_01_ | -0.534 |  |  |  |  |
|  | -0.439 |  |  |  |  |
|  | 0.177 |  |  |  |  |
|  | 0.095 |  |  |  |  |
| ICC | 0.886 |  |  |  |  |
| N _id_ | 128 |  |  |  |  |
| Observations | 1280 | | | | |
| Marginal R^2^ / Conditional R^2^ | 0.023 / 0.889 | | | | |

*Note.* The table shows the regression coefficients of fixed effects and summary information of random effects. The marginal R^2^ considers only the variance of fixed effects, while the conditional R^2^ takes both fixed and random effects into account. Degrees of freedom are based on Satterthwaite’s approximation. Statistically significant terms are highlighted in bold. Terms in brackets indicate the level of factor that is contrasted against the reference level, which is Epoch 1 for the Epoch factor and high-probability for the Triplet type factor.

### Model_SL-LDAR_

**Supplementary Table S6.** Contrasts by epoch between the estimates of LDAR.

| **Contrast** | ***p*** |
| --- | --- |
| Epoch 0 - Epoch 1 | **<0.001** |
| Epoch 0 - Epoch 2 | **<0.001** |
| Epoch 0 - Epoch 3 | **<0.001** |
| Epoch 0 - Epoch 4 | **<0.001** |
| Epoch 1 - Epoch 2 | 0.282 |
| Epoch 1 - Epoch 3 | **0.033** |
| Epoch 1 - Epoch 4 | **<0.001** |
| Epoch 2 - Epoch 3 | 0.996 |
| Epoch 2 - Epoch 4 | 0.114 |
| Epoch 3 - Epoch 4 | 0.598 |

**Supplementary Table S7.** Results of the linear mixed model predicting epochwise mean LDAR.

| **Prediction of LDAR** | | | | | |
| --- | --- | --- | --- | --- | --- |
| **Fixed Effects** | ***β estimates*** | ***95% CI*** | ***t*** | ***df*** | ***p*** |
| **(Intercept)** | 0.292 | 0.286 – 0.298 | 97.103 | 127.000 | **<0.001** |
| **Epoch [1]** | -0.041 | -0.047 – -0.035 | -13.130 | 508.000 | **<0.001** |
| Epoch [2] | -0.002 | -0.008 – 0.004 | -0.577 | 508.000 | 0.564 |
| **Epoch [3]** | 0.009 | 0.003 – 0.015 | 2.814 | 508.000 | **0.005** |
| **Epoch [4]** | 0.013 | 0.007 – 0.019 | 4.091 | 508.000 | **<0.001** |
| **Random Effects** | | | | | |
| σ^2^ | 0.002 |  |  |  |  |
| τ_00_ _id_ | 0.001 |  |  |  |  |
| ICC | 0.350 |  |  |  |  |
| N _id_ | 128 |  |  |  |  |
| Observations | 640 |  |  |  |  |
| Marginal R^2^ / Conditional R^2^ | 0.165 / 0.458 |  |  |  |  |

*Note.* The table shows the regression coefficients of fixed effects and summary information of random effects. The marginal R^2^ considers only the variance of fixed effects, while the conditional R^2^ takes both fixed and random effects into account. Degrees of freedom are based on Satterthwaite’s approximation. Statistically significant terms are highlighted in bold. Terms in brackets indicate the level of factor that is contrasted against the reference level, which is Epoch 1 for the Epoch factor.

### Model_saccade-LLH_

**Supplementary Table S8.** Estimates for the epochwise mean likelihood of saccade types.

| **Epoch** | **Type** | ***Estimated Mean*** | ***SE*** | ***df*** | ***95% CI*** |
| --- | --- | --- | --- | --- | --- |
| 0 | LD correct | 1.015 | 0.0198 | 2540 | [0.976, 1.054] |
| 0 | LD error | 1.001 | 0.0198 | 2540 | [0.962, 1.040] |
| 0 | NLD correct | 0.993 | 0.0198 | 2540 | [0.954, 1.032] |
| 0 | NLD error | 1.000 | 0.0198 | 2540 | [0.962, 1.039] |
| 1 | LD correct | 1.102 | 0.0198 | 2540 | [1.064, 1.141] |
| 1 | LD error | 1.243 | 0.0198 | 2540 | [1.204, 1.282] |
| 1 | NLD correct | 0.969 | 0.0198 | 2540 | [0.930, 1.007] |
| 1 | NLD error | 0.945 | 0.0198 | 2540 | [0.907, 0.984] |
| 2 | LD correct | 1.138 | 0.0198 | 2540 | [1.100, 1.177] |
| 2 | LD error | 1.304 | 0.0198 | 2540 | [1.266, 1.343] |
| 2 | NLD correct | 0.935 | 0.0198 | 2540 | [0.896, 0.974] |
| 2 | NLD error | 0.933 | 0.0198 | 2540 | [0.894, 0.972] |
| 3 | LD correct | 1.163 | 0.0198 | 2540 | [1.124, 1.201] |
| 3 | LD error | 1.312 | 0.0198 | 2540 | [1.273, 1.351] |
| 3 | NLD correct | 0.928 | 0.0198 | 2540 | [0.889, 0.967] |
| 3 | NLD error | 0.927 | 0.0198 | 2540 | [0.888, 0.966] |
| 4 | LD correct | 1.201 | 0.0198 | 2540 | [1.162, 1.239] |
| 4 | LD error | 1.333 | 0.0198 | 2540 | [1.294, 1.372] |
| 4 | NLD correct | 0.918 | 0.0198 | 2540 | [0.879, 0.957] |
| 4 | NLD error | 0.916 | 0.0198 | 2540 | [0.878, 0.955] |

**Supplementary Table S9.** Contrasts by epoch between estimates for the epochwise mean likelihood of saccade types.

| **Epoch** | **Contrast** | ***p*** |
| --- | --- | --- |
| 0 | LD correct - LD error | 0.997 |
| 0 | LD correct - NLD correct | 0.966 |
| 0 | LD correct - NLD error | 0.996 |
| 0 | LD error - NLD correct | 0.999 |
| 0 | LD error - NLD error | 1.000 |
| 0 | NLD correct - NLD error | 0.999 |
| 1 | LD correct - LD error | **<0.001** |
| 1 | LD correct - NLD correct | **<0.001** |
| 1 | LD correct - NLD error | **<0.001** |
| 1 | LD error - NLD correct | **<0.001** |
| 1 | LD error - NLD error | **<0.001** |
| 1 | NLD correct - NLD error | 0.957 |
| 2 | LD correct - LD error | **<0.001** |
| 2 | LD correct - NLD correct | **<0.001** |
| 2 | LD correct - NLD error | **<0.001** |
| 2 | LD error - NLD correct | **<0.001** |
| 2 | LD error - NLD error | **<0.001** |
| 2 | NLD correct - NLD error | 1.000 |
| 3 | LD correct - LD error | **<0.001** |
| 3 | LD correct - NLD correct | **<0.001** |
| 3 | LD correct - NLD error | **<0.001** |
| 3 | LD error - NLD correct | **<0.001** |
| 3 | LD error - NLD error | **<0.001** |
| 3 | NLD correct - NLD error | 1.000 |
| 4 | LD correct - LD error | **<0.001** |
| 4 | LD correct - NLD correct | **<0.001** |
| 4 | LD correct - NLD error | **<0.001** |
| 4 | LD error - NLD correct | **<0.001** |
| 4 | LD error - NLD error | **<0.001** |
| 4 | NLD correct - NLD error | 1.000 |

**Supplementary Table S10.** Contrasts by saccade type between estimates for the epochwise mean likelihood of saccade types.

| **Type** | **Contrast** | ***p*** |
| --- | --- | --- |
| LD correct | Epoch 0 - Epoch 1 | **0.017** |
| LD correct | Epoch 0 - Epoch 2 | **0.001** |
| LD correct | Epoch 0 - Epoch 3 | **< 0.001** |
| LD correct | Epoch 0 - Epoch 4 | **<0.001** |
| LD correct | Epoch 1 - Epoch 2 | 0.892 |
| LD correct | Epoch 1 - Epoch 3 | 0.273 |
| LD correct | Epoch 1 - Epoch 4 | **0.045** |
| LD correct | Epoch 2 - Epoch 3 | 0.992 |
| LD correct | Epoch 2 - Epoch 4 | 0.232 |
| LD correct | Epoch 3 - Epoch 4 | 0.854 |
| LD error | Epoch 0 - Epoch 1 | **<0.001** |
| LD error | Epoch 0 - Epoch 2 | **<0.001** |
| LD error | Epoch 0 - Epoch 3 | **< 0.001** |
| LD error | Epoch 0 - Epoch 4 | **<0.001** |
| LD error | Epoch 1 - Epoch 2 | 0.252 |
| LD error | Epoch 1 - Epoch 3 | 0.130 |
| LD error | Epoch 1 - Epoch 4 | **0.013** |
| LD error | Epoch 2 - Epoch 3 | 1.000 |
| LD error | Epoch 2 - Epoch 4 | 0.973 |
| LD error | Epoch 3 - Epoch 4 | 0.997 |
| NLD correct | Epoch 0 - Epoch 1 | 0.992 |
| NLD correct | Epoch 0 - Epoch 2 | 0.323 |
| NLD correct | Epoch 0 - Epoch 3 | 0.185 |
| NLD correct | Epoch 0 - Epoch 4 | 0.072 |
| NLD correct | Epoch 1 - Epoch 2 | 0.926 |
| NLD correct | Epoch 1 - Epoch 3 | 0.795 |
| NLD correct | Epoch 1 - Epoch 4 | 0.521 |
| NLD correct | Epoch 2 - Epoch 3 | 1.000 |
| NLD correct | Epoch 2 - Epoch 4 | 0.999 |
| NLD correct | Epoch 3 - Epoch 4 | 1.000 |
| NLD error | Epoch 0 - Epoch 1 | 0.395 |
| NLD error | Epoch 0 - Epoch 2 | 0.147 |
| NLD error | Epoch 0 - Epoch 3 | 0.083 |
| NLD error | Epoch 0 - Epoch 4 | **0.026** |
| NLD error | Epoch 1 - Epoch 2 | 1.000 |
| NLD error | Epoch 1 - Epoch 3 | 0.999 |
| NLD error | Epoch 1 - Epoch 4 | 0.972 |
| NLD error | Epoch 2 - Epoch 3 | 1.000 |
| NLD error | Epoch 2 - Epoch 4 | 0.999 |
| NLD error | Epoch 3 - Epoch 4 | 1.000 |

**Supplementary Table S11.** Results of the linear mixed model predicting the epochwise likelihood of saccade types.

| **Prediction of saccade type likelihood** | | | | | |
| --- | --- | --- | --- | --- | --- |
| **Fixed Effects** | ***β estimates*** | ***95% CI*** | ***t*** | ***df*** | ***p*** |
| **(Intercept)** | 1.064 | 1.055 – 1.073 | 233.808 | 127 | **<0.001** |
| **Type [LD correct]** | 0.060 | 0.045 – 0.075 | 7.831 | 2413 | **<0.001** |
| **Type [LD error]** | 0.175 | 0.160 – 0.190 | 22.861 | 2413 | **<0.001** |
| **Type [NLD correct]** | -0.115 | -0.130 – -0.100 | -15.081 | 2413 | **<0.001** |
| Epoch [1] | -0.062 | -0.079 – -0.044 | -6.970 | 2413 | **<0.001** |
| Epoch [2] | 0.001 | -0.016 – 0.018 | 0.118 | 2413 | 0.906 |
| Epoch [3] | 0.014 | -0.004 – 0.031 | 1.560 | 2413 | 0.119 |
| Epoch [4] | 0.019 | 0.001 – 0.036 | 2.101 | 2413 | **0.036** |
| **Type [LD correct] × Epoch [1]** | -0.047 | -0.077 – -0.017 | -3.097 | 2413 | **0.002** |
| **Type [LD error] × Epoch [1]** | -0.176 | -0.206 – -0.146 | -11.511 | 2413 | **<0.001** |
| **Type [NLD correct] × Epoch [1]** | 0.106 | 0.076 – 0.136 | 6.924 | 2413 | **<0.001** |
| Type [LD correct] × Epoch [2] | -0.022 | -0.052 – 0.008 | -1.459 | 2413 | 0.145 |
| Type [LD error] × Epoch [2] | 0.003 | -0.027 – 0.033 | 0.221 | 2413 | 0.825 |
| Type [NLD correct] × Epoch [2] | 0.019 | -0.011 – 0.049 | 1.243 | 2413 | 0.214 |
| Type [LD correct] × Epoch [3] | 0.001 | -0.029 – 0.031 | 0.052 | 2413 | 0.959 |
| Type [LD error] × Epoch [3] | 0.052 | 0.022 – 0.082 | 3.387 | 2413 | **0.001** |
| Type [NLD correct] × Epoch [3] | -0.027 | -0.057 – 0.003 | -1.785 | 2413 | 0.074 |
| Type [LD correct] × Epoch [4] | 0.020 | -0.010 – 0.050 | 1.328 | 2413 | 0.184 |
| Type [LD error] × Epoch [4] | 0.055 | 0.025 – 0.085 | 3.575 | 2413 | **<0.001** |
| Type [NLD correct] × Epoch [4] | -0.039 | -0.069 – -0.009 | -2.553 | 2413 | **0.011** |
| **Random Effects** | | | | | |
| σ^2^ | 0.050 |  |  |  |  |
| τ_00_ _id_ | 0.000 |  |  |  |  |
| ICC | 0.003 |  |  |  |  |
| N _id_ | 128 |  |  |  |  |
| Observations | 2560 |  |  |  |  |
| Marginal R^2^ / Conditional R^2^ | 0.291 / 0.294 |  |  |  |  |

*Note.* The table shows the regression coefficients of fixed effects and summary information of random effects. The marginal R^2^ considers only the variance of fixed effects, while the conditional R^2^ takes both fixed and random effects into account. Degrees of freedom are based on Satterthwaite’s approximation. Statistically significant terms are highlighted in bold. Terms in brackets indicate the level of factor that is contrasted against the reference level, which is Epoch 0 for the Epoch factor and NLD error for the Type factor.

### Raw number and percentage of trials

**Supplementary Table S12.** Epochwise number and percentage of updates and no updates.

| **Epoch** | Update | No update | Total |
| --- | --- | --- | --- |
| **1** | 14695 (60.30%) | 9674 (39.70%) | 24369 |
| **2** | 14955 (60.49%) | 9769 (39.51%) | 24724 |
| **3** | 14792 (60.10%) | 9822 (39.90%) | 24614 |
| **4** | 14990 (60.28%) | 9878 (39.72%) | 24868 |

### Model_saccade-update_

**Supplementary Table S13.** Estimates for the mean likelihood of saccade types.

| **Type** | ***Probability*** | ***SE*** | ***95% CI*** |
| --- | --- | --- | --- |
| LD correct | 0.582 | 0.0074 | [0.567, 0.596] |
| LD error | 0.568 | 0.0074 | [0.553, 0.584] |
| NLD correct | 0.640 | 0.0074 | [0.624, 0.655] |
| NLD error | 0.630 | 0.0074 | [0.617, 0.642] |

**Supplementary Table S14.** Contrasts for the mean likelihood of saccade types.

| **Contrast** | ***p*** |
| --- | --- |
| LD correct - LD error | 0.133 |
| LD correct - NLD correct | **<0.001** |
| LD correct - NLD error | **<0.001** |
| LD error - NLD correct | **<0.001** |
| LD error - NLD error | **<0.001** |
| NLD correct - NLD error | 0.352 |

**Supplementary Table S15.** Results of the generalized linear mixed model predicting trialwise update.

| **Prediction of trialwise update** | | | | | |
| --- | --- | --- | --- | --- | --- |
| **Fixed Effects** | ***Odds Ratios*** | ***95% CI*** | ***z*** | ***df*** | ***p*** |
| **(Intercept)** | 0.425 | 0.338 – 0.533 | -7.365 | Inf | **<0.001** |
| **Epoch [2]** | 1.057 | 1.019 – 1.096 | 2.940 | Inf | **0.003** |
| Epoch [3] | 1.034 | 1.000 – 1.069 | 1.958 | Inf | 0.050 |
| Epoch [4] | 0.982 | 0.950 – 1.016 | -1.049 | Inf | 0.294 |
| **Previous saccade type [LD correct]** | 1.079 | 1.045 – 1.113 | 4.748 | Inf | **<0.001** |
| **Previous saccade type [LD error]** | 0.873 | 0.841 – 0.907 | -7.029 | Inf | **<0.001** |
| **Previous saccade type [NLD correct]** | 1.071 | 1.028 – 1.117 | 3.280 | Inf | **0.001** |
| Epoch [2] × Previous saccade type [LD correct] | 1.042 | 0.985 – 1.102 | 1.422 | Inf | 0.155 |
| Epoch [3] × Previous saccade type [LD correct] | 1.000 | 0.948 – 1.055 | 0.005 | Inf | 0.996 |
| Epoch [4] × Previous saccade type [LD correct] | 0.960 | 0.910 – 1.012 | -1.518 | Inf | 0.129 |
| Epoch [2] × Previous saccade type [LD error] | 1.050 | 0.977 – 1.129 | 1.331 | Inf | 0.183 |
| **Epoch [3] × Previous saccade type [LD error]** | 0.929 | 0.872 – 0.990 | -2.270 | Inf | **0.023** |
| Epoch [4] × Previous saccade type [LD error] | 1.040 | 0.977 – 1.108 | 1.240 | Inf | 0.215 |
| Epoch [2] × Previous saccade type [NLD correct] | 0.934 | 0.864 – 1.009 | -1.736 | Inf | 0.083 |
| Epoch [3] × Previous saccade type [NLD correct] | 1.070 | 0.999 – 1.146 | 1.920 | Inf | 0.055 |
| Epoch [4] × Previous saccade type [NLD correct] | 1.017 | 0.948 – 1.090 | 0.465 | Inf | 0.642 |
| **Random Effects** | | | | | |
| σ^2^ | 3.290 |  |  |  |  |
| τ_00_ _id_ | 0.846 |  |  |  |  |
| ICC | 0.205 |  |  |  |  |
| N _id_ | 61 |  |  |  |  |
| Observations | 93696 |  |  |  |  |
| Marginal R^2^ / Conditional R^2^ | 0.001 / 0.206 |  |  |  |  |

*Note.* The table shows the regression coefficients of fixed effects and summary information of random effects. P values from statistics are from z-tests, making the degrees of freedom infinite. R^2^ considers only the variance of fixed effects, while the conditional R^2^ takes both fixed and random effects into account. Statistically significant terms are highlighted in bold. Terms in brackets indicate the level of factor that is contrasted against the reference level, which is Epoch 1 for the Epoch factor and NLD error for the Previous saccade type factor.

**Supplementary Table S16.** Contrasts for the mean likelihood of saccade types per trials since triplet

| **Trials since triplet** | **Contrast** | ***p*** |
| --- | --- | --- |
| Mean - 1 SD | LD correct - LD error | 1.000 |
| Mean - 1 SD | LD correct - NLD correct | **<0.001** |
| Mean - 1 SD | LD correct - NLD error | **<0.001** |
| Mean - 1 SD | LD error - NLD correct | **<0.001** |
| Mean - 1 SD | LD error - NLD error | **<0.001** |
| Mean - 1 SD | NLD correct - NLD error | 1.000 |
| Mean + 1 SD | LD correct - LD error | **0.001** |
| Mean + 1 SD | LD correct - NLD correct | **<0.001** |
| Mean + 1 SD | LD correct - NLD error | **<0.001** |
| Mean + 1 SD | LD error - NLD correct | **<0.001** |
| Mean + 1 SD | LD error - NLD error | **<0.001** |
| Mean + 1 SD | NLD correct - NLD error | 0.999 |

**Supplementary Table S17.** Contrasts for the trials since triplet per saccade types

| **Type** | **Contrast** | ***p*** |
| --- | --- | --- |
| LD correct | Mean - 1 SD - Mean + 1 SD | 1.000 |
| LD error | Mean - 1 SD - Mean + 1 SD | 0.197 |
| NLD correct | Mean - 1 SD - Mean + 1 SD | **<0.001** |
| NLD error | Mean - 1 SD - Mean + 1 SD | **<0.001** |

**Supplementary Table S18.** Median trials since triplet per previous saccade type per epoch

| **Epoch** | **Type** | ***Median*** |
| --- | --- | --- |
| 1 | LD correct | 68 |
| 1 | LD error | 262 |
| 1 | NLD correct | 279 |
| 1 | NLD error | 105 |
| 2 | LD correct | 70 |
| 2 | LD error | 292 |
| 2 | NLD correct | 312 |
| 2 | NLD error | 107 |
| 3 | LD correct | 68 |
| 3 | LD error | 325 |
| 3 | NLD correct | 328 |
| 3 | NLD error | 104 |
| 4 | LD correct | 69 |
| 4 | LD error | 316 |
| 4 | NLD correct | 325 |
| 4 | NLD error | 106 |

**Supplementary Table S19.** Results of the generalized linear mixed model predicting trialwise update including the variable trials since the last occurrence of the triplet.

| **Prediction of trialwise update** | | | |
| --- | --- | --- | --- |
| ***Predictors*** | ***Odds Ratios*** | ***CI*** | ***p*** |
| **(Intercept)** | 1.53 | 1.45 – 1.62 | **<0.001** |
| Epoch | 0.99 | 0.97 – 1.01 | 0.234 |
| **Previous saccade type [LD correct]** | 0.95 | 0.92 – 0.99 | **0.009** |
| **Previous saccade type [LD error]** | 0.84 | 0.82 – 0.87 | **<0.001** |
| **Previous saccade type [NLD correct]** | 1.12 | 1.07 – 1.16 | **<0.001** |
| **Trials since triplet** | 1.08 | 1.06 – 1.11 | **<0.001** |
| Epoch × Previous saccade type [LD correct] | 1.00 | 0.96 – 1.03 | 0.939 |
| **Epoch × Previous saccade type [LD error]** | 0.94 | 0.91 – 0.98 | **0.001** |
| **Epoch × Previous saccade type [NLD correct]** | 1.04 | 1.00 – 1.08 | **0.032** |
| Epoch × Trials since triplet | 0.98 | 0.96 – 1.01 | 0.125 |
| **Previous saccade type [LD correct] × Trials since triplet** | 1.07 | 1.01 – 1.14 | **0.026** |
| **Previous saccade type [LD error] × Trials since triplet** | 0.94 | 0.91 – 0.97 | **<0.001** |
| Previous saccade type [NLD correct] × Trials since triplet | 1.00 | 0.96 – 1.03 | 0.929 |
| Epoch × Previous saccade type [LD correct] × Trial since triplet | 1.03 | 0.97 – 1.09 | 0.386 |
| Epoch × Previous saccade type [LD error] × Trials since triplet | 1.03 | 1.00 – 1.07 | 0.068 |
| **Epoch × Previous saccade type [NLD correct]× Trials since triplet** | 0.95 | 0.92 – 0.99 | **0.005** |
| **Random Effects** | | | |
| σ^2^ | 3.29 | | |
| τ_00_ _id_ | 0.08 | | |
| ICC | 0.03 | | |
| N _id_ | 128 | | |
| Observations | 98575 | | |
| Marginal R^2^ / Conditional R^2^ | 0.005 / 0.030 | | |

*Note.* The table shows the regression coefficients of fixed effects and summary information of random effects. Both Epoch and Trial since triplet was z-scored. R^2^ considers only the variance of fixed effects, while the conditional R^2^ takes both fixed and random effects into account. Statistically significant terms are highlighted in bold. Terms in brackets indicate the level of factor that is contrasted against the reference level, which is NLD error for the Previous saccade type factor.

**Supplementary Figure S1.** Predicted probability of update per previous saccade type.

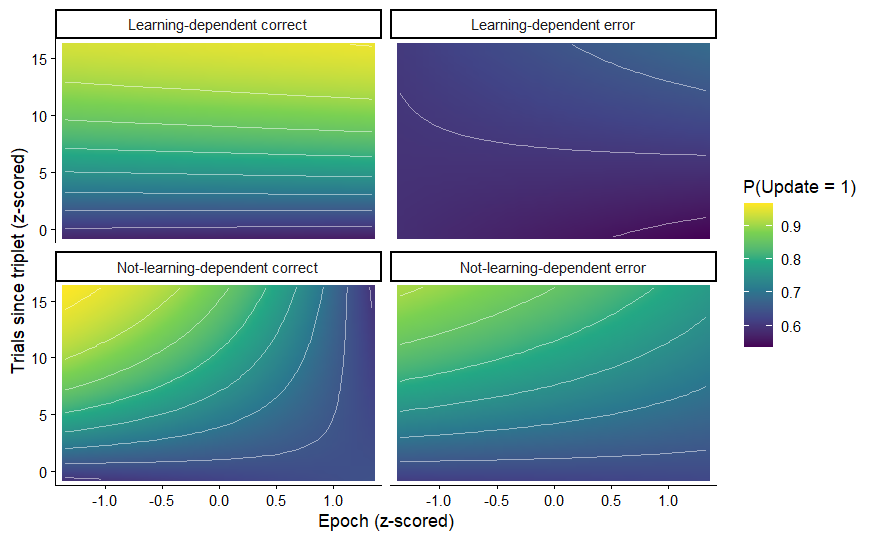

*Note.* The panels show model-predicted probabilities of an update from a generalized linear mixed-effects model. Predictions are plotted across the continuous ranges of z-scored epochs and trials since triplet, separately for each previous saccade type. Warmer colors indicate higher predicted probabilities of an update. Contour lines represent iso-probability levels. Surfaces illustrate how the probability of updating varies jointly as a function of learning stage and recency of triplet exposure, and how this relationship differs between saccade types.

### Raw number and percentage of trials

**Supplementary Table S20.** Epochwise number and percentage of trials in each update type

| **Epoch** | LD same | LD-to- NLD update | NLD same | NLD-to LD update | NLD-to- other-NLD update | Total |
| --- | --- | --- | --- | --- | --- | --- |
| **1** | 2862 (11.74%) | 3917 (16.07%) | 6812 (27.95%) | 4148 (17.02%) | 6630 (27.21%) | 24369 |
| **2** | 3136 (12.68%) | 4100 (16.58%) | 6633 (26.83%) | 4257 (17.22%) | 6598 (26.69%) | 24724 |
| **3** | 3257 (13.23%) | 4180 (16.98%) | 6565 (26.67%) | 4191 (07.03%) | 6421 (26.09%) | 24614 |
| **4** | 3435 (13.81%) | 4312 (17.34%) | 6443 (25.91%) | 4327 (17.40%) | 6351 (25.54%) | 24868 |

### Model_update-LLH_

**Supplementary Table S21.** Estimates for the epochwise mean likelihood of update types.

| **Epoch** | **Type** | ***Estimated Mean*** | ***SE*** | ***df*** | ***95% CI*** |
| --- | --- | --- | --- | --- | --- |
| 1 | LD same | 1.840 | 0.0332 | 2538 | [1.775, 1.905] |
| 1 | LD-to-NLD | 0.870 | 0.0332 | 2538 | [0.805, 0.935] |
| 1 | NLD same | 1.441 | 0.0332 | 2538 | [1.376, 1.506] |
| 1 | NLD-to-LD | 0.934 | 0.0332 | 2538 | [0.869, 0.999] |
| 1 | NLD-to-other-NLD | 0.738 | 0.0332 | 2538 | [0.673, 0.803] |
| 2 | LD same | 1.993 | 0.0332 | 2538 | [1.927, 2.058] |
| 2 | LD-to-NLD | 0.896 | 0.0332 | 2538 | [0.831, 0.961] |
| 2 | NLD same | 1.386 | 0.0332 | 2538 | [1.321, 1.451] |
| 2 | NLD-to-LD | 0.935 | 0.0332 | 2538 | [0.870, 1.000] |
| 2 | NLD-to-other-NLD | 0.726 | 0.0332 | 2538 | [0.661, 0.791] |
| 3 | LD same | 2.101 | 0.0332 | 2538 | [2.035, 2.166] |
| 3 | LD-to-NLD | 0.921 | 0.0332 | 2538 | [0.856, 0.987] |
| 3 | NLD same | 1.363 | 0.0332 | 2538 | [1.298, 1.428] |
| 3 | NLD-to-LD | 0.924 | 0.0332 | 2538 | [0.859, 0.989] |
| 3 | NLD-to-other-NLD | 0.712 | 0.0332 | 2538 | [0.647, 0.778] |
| 4 | LD same | 2.170 | 0.0332 | 2538 | [2.105, 2.235] |
| 4 | LD-to-NLD | 0.934 | 0.0332 | 2538 | [0.869, 0.999] |
| 4 | NLD same | 1.351 | 0.0332 | 2538 | [1.286, 1.417] |
| 4 | NLD-to-LD | 0.944 | 0.0332 | 2538 | [0.879, 1.009] |
| 4 | NLD-to-other-NLD | 0.690 | 0.0332 | 2538 | [0.625, 0.755] |

**Supplementary Table S22.** Contrasts by epoch between estimates for the epochwise mean ratio of update types.

| **Epoch** | **Contrast** | ***p*** |
| --- | --- | --- |
| 1 | LD same - LD-to-NLD | **<0.001** |
| 1 | LD same - NLD same | **<0.001** |
| 1 | LD same - NLD-to-LD | **<0.001** |
| 1 | LD same - NLD-to-other-NLD | **<0.001** |
| 1 | LD-to-NLD - NLD same | **<0.001** |
| 1 | LD-to-NLD - NLD-to-LD | 0.854 |
| 1 | LD-to-NLD - NLD-to-other-NLD | **0.046** |
| 1 | NLD-same - NLD-to-LD | **<0.001** |
| 1 | NLD same - NLD-to-other-NLD | **<0.001** |
| 1 | NLD-to-LD - NLD-to-other-NLD | **0.003** |
| 2 | LD same - LD-to-NLD | **<0.001** |
| 2 | LD same - NLD same | **<0.001** |
| 2 | LD same - NLD-to-LD | **<0.001** |
| 2 | LD same - NLD-to-other-NLD | **<0.001** |
| 2 | LD-to-NLD - NLD same | **<0.001** |
| 2 | LD-to-NLD - NLD-to-LD | 0.994 |
| 2 | LD-to-NLD - NLD-to-other-NLD | **0.003** |
| 2 | NLD-same - NLD-to-LD | **<0.001** |
| 2 | NLD same - NLD-to-other-NLD | **<0.001** |
| 2 | NLD-to-LD - NLD-to-other-NLD | **0.001** |
| 3 | LD same - LD-to-NLD | **<0.001** |
| 3 | LD same - NLD same | **<0.001** |
| 3 | LD same - NLD-to-LD | **<0.001** |
| 3 | LD same - NLD-to-other-NLD | **<0.001** |
| 3 | LD-to-NLD - NLD same | **<0.001** |
| 3 | LD-to-NLD - NLD-to-LD | 1.000 |
| 3 | LD-to-NLD - NLD-to-other-NLD | **0.001** |
| 3 | NLD-same - NLD-to-LD | **<0.001** |
| 3 | NLD same - NLD-to-other-NLD | **<0.001** |
| 3 | NLD-to-LD - NLD-to-other-NLD | **0.001** |
| 4 | LD same - LD-to-NLD | **<0.001** |
| 4 | LD same - NLD same | **<0.001** |
| 4 | LD same - NLD-to-LD | **<0.001** |
| 4 | LD same - NLD-to-other-NLD | **<0.001** |
| 4 | LD-to-NLD - NLD same | **<0.001** |
| 4 | LD-to-NLD - NLD-to-LD | 1.000 |
| 4 | LD-to-NLD - NLD-to-other-NLD | **0.001** |
| 4 | NLD-same - NLD-to-LD | **<0.001** |
| 4 | NLD same - NLD-to-other-NLD | **<0.001** |
| 4 | NLD-to-LD - NLD-to-other-NLD | **<0.001** |

**Supplementary Table S23.** Contrasts by saccade type between estimates for the epochwise mean ratio of update types.

| **Type** | **Contrast** | ***p*** |
| --- | --- | --- |
| LD same | Epoch 1 - Epoch 2 | 0.067 |
| LD same | Epoch 1 - Epoch 3 | **<0.001** |
| LD same | Epoch 1 - Epoch 4 | **<0.001** |
| LD same | Epoch 2 - Epoch 3 | 0.120 |
| LD same | Epoch 2 - Epoch 4 | **0.009** |
| LD same | Epoch 3 - Epoch 4 | 0.593 |
| LD-to-NLD | Epoch 1 - Epoch 2 | 0.995 |
| LD-to-NLD | Epoch 1 - Epoch 3 | 0.852 |
| LD-to-NLD | Epoch 1 - Epoch 4 | 0.679 |
| LD-to-NLD | Epoch 2 - Epoch 3 | 0.995 |
| LD-to-NLD | Epoch 2 - Epoch 4 | 0.959 |
| LD-to-NLD | Epoch 3 - Epoch 4 | 0.999 |
| NLD same | Epoch 1 - Epoch 2 | 0.805 |
| NLD same | Epoch 1 - Epoch 3 | 0.454 |
| NLD same | Epoch 1 - Epoch 4 | 0.291 |
| NLD same | Epoch 2 - Epoch 3 | 0.997 |
| NLD same | Epoch 2 - Epoch 4 | 0.976 |
| NLD same | Epoch 3 - Epoch 4 | 0.999 |
| NLD-to-LD | Epoch 1 - Epoch 2 | 1.000 |
| NLD-to-LD | Epoch 1 - Epoch 3 | 1.000 |
| NLD-to-LD | Epoch 1 - Epoch 4 | 1.000 |
| NLD-to-LD | Epoch 2 - Epoch 3 | 1.000 |
| NLD-to-LD | Epoch 2 - Epoch 4 | 1.000 |
| NLD-to-LD | Epoch 3 - Epoch 4 | 0.999 |
| NLD-to-other-NLD | Epoch 1 - Epoch 2 | 1.000 |
| NLD-to-other-NLD | Epoch 1 - Epoch 3 | 0.995 |
| NLD-to-other-NLD | Epoch 1 - Epoch 4 | 0.895 |
| NLD-to-other-NLD | Epoch 2 - Epoch 3 | 0.999 |
| NLD-to-other-NLD | Epoch 2 - Epoch 4 | 0.970 |
| NLD-to-other-NLD | Epoch 3 - Epoch 4 | 0.998 |

**Supplementary Table S24.** Results of the linear mixed model predicting the epochwise likelihood of update types.

| **Prediction of update type likelihood** | | | | | |
| --- | --- | --- | --- | --- | --- |
| **Fixed Effects** | ***β estimates*** | ***95% CI*** | ***t*** | ***df*** | ***p*** |
| **(Intercept)** | 1.193 | 1.178 – 1.209 | 152.241 | 59.000 | **<0.001** |
| **LD same** | 0.832 | 0.803 – 0.861 | 56.239 | 1121.000 | **<0.001** |
| **LD-to-NLD** | -0.288 | -0.317 – -0.259 | -19.466 | 1121.000 | **<0.001** |
| **NLD-same** | 0.192 | 0.163 – 0.221 | 12.970 | 1121.000 | **<0.001** |
| **NLD-to-LD** | -0.259 | -0.288 – -0.230 | -17.527 | 1121.000 | **<0.001** |
| **Epoch [2]** | -0.029 | -0.054 – 0.004 | -2.260 | 1121.000 | **0.024** |
| Epoch [3] | -0.006 | -0.031 – 0.019 | -0.494 | 1121.000 | 0.621 |
| Epoch [4] | 0.011 | -0.014 – 0.036 | 0.845 | 1121.000 | 0.398 |
| **LD same × Epoch [2]** | -0.157 | -0.207– -0.107 | -6.121 | 1121.000 | **<0.001** |
| LD-to-NLD × Epoch [2] | -0.006 | -0.056 – 0.044 | -0.243 | 1121.000 | 0.808 |
| **NLD-same × Epoch [2]** | 0.085 | 0.034 – 0.135 | 3.302 | 1121.000 | **0.001** |
| NLD-to-LD × Epoch [2] | 0.029 | -0.022 – 0.079 | 1.115 | 1121.000 | 0.265 |
| LD same × Epoch [3] | -0.027 | -0.077 – 0.023 | -1.045 | 1121.000 | 0.296 |
| LD-to-NLD × Epoch [3] | -0.003 | -0.054 – 0.047 | -0.128 | 1121.000 | 0.898 |
| NLD-same × Epoch [3] | 0.007 | -0.043 – 0.057 | 0.268 | 1121.000 | 0.789 |
| NLD-to-LD × Epoch [3] | 0.007 | -0.043 – 0.058 | 0.284 | 1121.000 | 0.777 |
| **LD same × Epoch [4]** | 0.064 | 0.014 – 0.114 | 2.499 | 1121.000 | **0.013** |
| LD-to-NLD × Epoch [4] | 0.005 | -0.045 – 0.056 | 0.207 | 1121.000 | 0.836 |
| NLD-same × Epoch [4] | -0.033 | -0.083 – 0.017 | -0.292 | 1121.000 | 0.197 |
| NLD-to-LD × Epoch [4] | -0.021 | -0.071 – 0.029 | -0.827 | 1121.000 | 0.408 |
| **Random Effects** | | | | | |
| σ^2^ | 0.140 |  |  |  |  |
| τ_00_ _id_ | 0.001 |  |  |  |  |
| ICC | 0.006 |  |  |  |  |
| N _id_ | 128 |  |  |  |  |
| Observations | 2560 |  |  |  |  |
| Marginal R^2^ / Conditional R^2^ | 0.615 / 0.617 |  |  |  |  |

*Note.* The table shows the regression coefficients of fixed effects and summary information of random effects. The marginal R^2^ considers only the variance of fixed effects, while the conditional R^2^ takes both fixed and random effects into account. Degrees of freedom are based on Satterthwaite’s approximation. Statistically significant terms are highlighted in bold. Terms in brackets indicate the level of factor that is contrasted against the reference level, which is Epoch 1 for the Epoch factor and NLD-to-other-NLD for the Type factor.
